# Proneural genes define ground state rules to regulate neurogenic patterning and cortical folding

**DOI:** 10.1101/2020.09.22.307058

**Authors:** Sisu Han, Grey A Wilkinson, Satoshi Okawa, Lata Adnani, Rajiv Dixit, Imrul Faisal, Matthew Brooks, Veronique Cortay, Vorapin Chinchalongporn, Dawn Zinyk, Saiqun Li, Jinghua Gao, Faizan Malik, Yacine Touahri, Vladimir Espinosa Angarica, Ana-Maria Oproescu, Eko Raharjo, Yaroslav Ilnytskyy, Jung-Woong Kim, Wei Wu, Waleed Rahmani, Igor Kovalchuk, Jennifer Ai-wen Chan, Deborah Kurrasch, Diogo S. Castro, Colette Dehay, Anand Swaroop, Jeff Biernaskie, Antonio del Sol, Carol Schuurmans

## Abstract

Transition from smooth, lissencephalic brains to highly-folded, gyrencephalic structures is associated with neuronal expansion and breaks in neurogenic symmetry. Here we show that *Neurog2* and *Ascl1* proneural genes regulate cortical progenitor cell differentiation through cross-repressive interactions to sustain neurogenic continuity in a lissencephalic rodent brain. Using *in vivo* lineage tracing, we found that *Neurog2* and *Ascl1* expression defines a lineage continuum of four progenitor pools, with ‘double^+^ progenitors’ displaying several unique features (least lineage-restricted, complex gene regulatory network, G_2_ pausing). Strikingly, selective killing of double^+^ progenitors using split-Cre;*Rosa-DTA* transgenics breaks neurogenic symmetry by locally disrupting Notch signaling, leading to cortical folding. Finally, consistent with *NEUROG2* and *ASCL1* driving discontinuous neurogenesis and folding in gyrencephalic species, their transcripts are modular in folded macaque cortices and pseudo-folded human cerebral organoids. *Neurog2*/*Ascl1* double^+^ progenitors are thus Notch-ligand expressing ‘niche’ cells that control neurogenic periodicity to determine cortical gyrification.

**HIGHLIGHTS:**

- Neurog2 and Ascl1 expression defines four distinct transitional progenitor states
- Double^+^ NPCs are transcriptionally complex and mark a lineage branch point
- Double^+^ NPCs control neurogenic patterning and cortical folding via Notch signaling
- Neurog2 and Ascl1 expression is modular in folded and not lissencephalic cortices

**eTOC BLURB:** Emergence of a gyrencephalic cortex is associated with a break in neurogenic continuity across the cortical germinal zone. Han et al. identify a pool of unbiased neural progenitors at a lineage bifurcation point that co-express Neurog2 and Ascl1 and produce Notch ligands to control neurogenic periodicity and cortical folding.

## INTRODUCTION

The cerebral cortex displays enormous structural diversity, transitioning during evolution between a highly folded, gyrencephalic structure in primates and larger mammals, to a smooth, lissencephalic form in smaller mammals (Lewitus et al., 2014). Despite this phenotypic diversity, the underlying genetic programs that guide cortical development are highly conserved (Kriegstein et al., 2006), raising the question of how built-in constraints have been modified to allow new cortical patterns to emerge. The blueprints for lissencephalic and gyrencephalic cortices are beginning to be deciphered, revealing a diversification of the central ‘building blocks’ – the multipotent neural stem (NSC) and progenitor (NPC, hereafter used collectively) cells that give rise to cortical neurons and glia. Cortical NPCs are stratified into apical and basal pools, the latter showing greater species divergence. In lissencephalic species, basal NPCs primarily include intermediate progenitor cells (IPCs), which have a limited proliferative capacity (Miyata et al., 2004). In gyrencephalic species, the basal NPC pool has expanded to form an outer subventricular zone (oSVZ) not present in rodents that contains outer radial glia (oRG) (Reillo et al., 2011). oRG give rise to transit-amplifying IPCs that divide several more times in gyrencephalic species to generate an increased number of later-born, upper-layer neurons that become folded (Molnar et al., 2011).

Genetic manipulations that asymmetrically expand the basal NPC pool can induce cortical folding in mice (Florio et al., 2015; Stahl et al., 2013), but oRG expansion alone does not explain why primary gyri (outward folds) and sulci (inward fissures) form in stereotyped positions, a constancy of phenotype that must be developmentally regulated. Local NPC expansion is unlikely to trigger folding as ‘inner’ germinal zones are ‘smooth’, and only outer neuronal layers are folded in gyrencephalic brains (Llinares-Benadero and Borrell, 2019). Instead, gyri develop in regions of high neurogenesis that are interspersed with regions of lower neurogenesis, where sulci form. Understanding how these neurogenic patterns arise requires new insights into how NPCs make the all-important decision to proliferate or differentiate.

Single cell transcriptomic analyses suggest that cortical NPCs undergo a continuum of state transitions before terminal differentiation (Telley et al., 2019), akin to the gradual acquisition of lineage bias in hematopoiesis (Brand and Morrissey, 2020). In hematopoiesis, lineage specifying transcription factors (TFs) and alterations to the chromatin landscape together control cell fate (Brand and Morrissey, 2020). In the cortex, proneural basic-helix-loop-helix (bHLH) TFs promote NPC differentiation and specify cell identities (Wilkinson et al., 2013), but how proneural gene expression/function relates to NPC transition states and changing fate probabilities is poorly understood. Three proneural TFs are expressed in cortical NPCs, *Neurog1* and *Neurog2*, which specify a glutamatergic neuronal identity, with *Neurog2* the dominant factor (Han et al., 2018), and *Ascl1*, the function of which is poorly understood in cortical NPCs (Wilkinson et al., 2013). Here we found that *Neurog2* and *Ascl1* expression biases cortical NPCs toward neuronal and oligodendrocyte fates, respectively, while double^+^ NPCs are less lineage restricted. Mechanistically, we show that *Neurog2* and *Ascl1* are cross repressive, thereby preventing lineage commitment by double^+^ NPCs. At the molecular level, *Neurog2* and *Ascl1* expression defines four NPC states defined by distinct fate probabilities, unique transcriptomes and gene regulatory networks, and a continuum of chromatin accessibility (double^+^>Ascl1^+^>proneural negative>Neurog2^+^). In sum, double^+^ NPCs define a lineage branch point, but also function as cortical niche cells that produce high levels of Notch ligands to modulate neurogenic patterning and control cortical folding.

## RESULTS

### Neurog2 and Ascl1 single^+^ NPCs are more lineage biased than double^+^ NPCs

Lissencephaly is associated with continuous neurogenesis across the cortical germinal zone, while gyrification evolved with symmetry-breaking neuronal expansion to form outward folds and inward fissures (Lewitus et al., 2014). Proneural genes are the main drivers of neurogenesis, prompting us to ask how they support a lissencephalic neurogenic pattern in mice. We focused on Neurog2 and Ascl1, as Neurog1 is almost exclusively expressed in Neurog2^+^ cortical NPCs and does not define a distinct NPC pool (Han et al., 2018). At E12.5, most DAPI^+^ NPCs in the cortical ventricular zone (VZ) did not express Neurog2 or Ascl1 and were called proneural negative (pro^−^) (78.64±1.31%, Figure 1A). Proneural positive (pro^+^) cells included Neurog2^+^ (15.07±1.46%), Ascl1^+^ (2.85±0.33%) and double^+^ (3.43±0.25%) NPCs (Figure 1A). The early cortical VZ can thus be stratified into four NPC pools: pro^−^, Neurog2^+^, Ascl1^+^ and double^+^.

**Figure 1.**
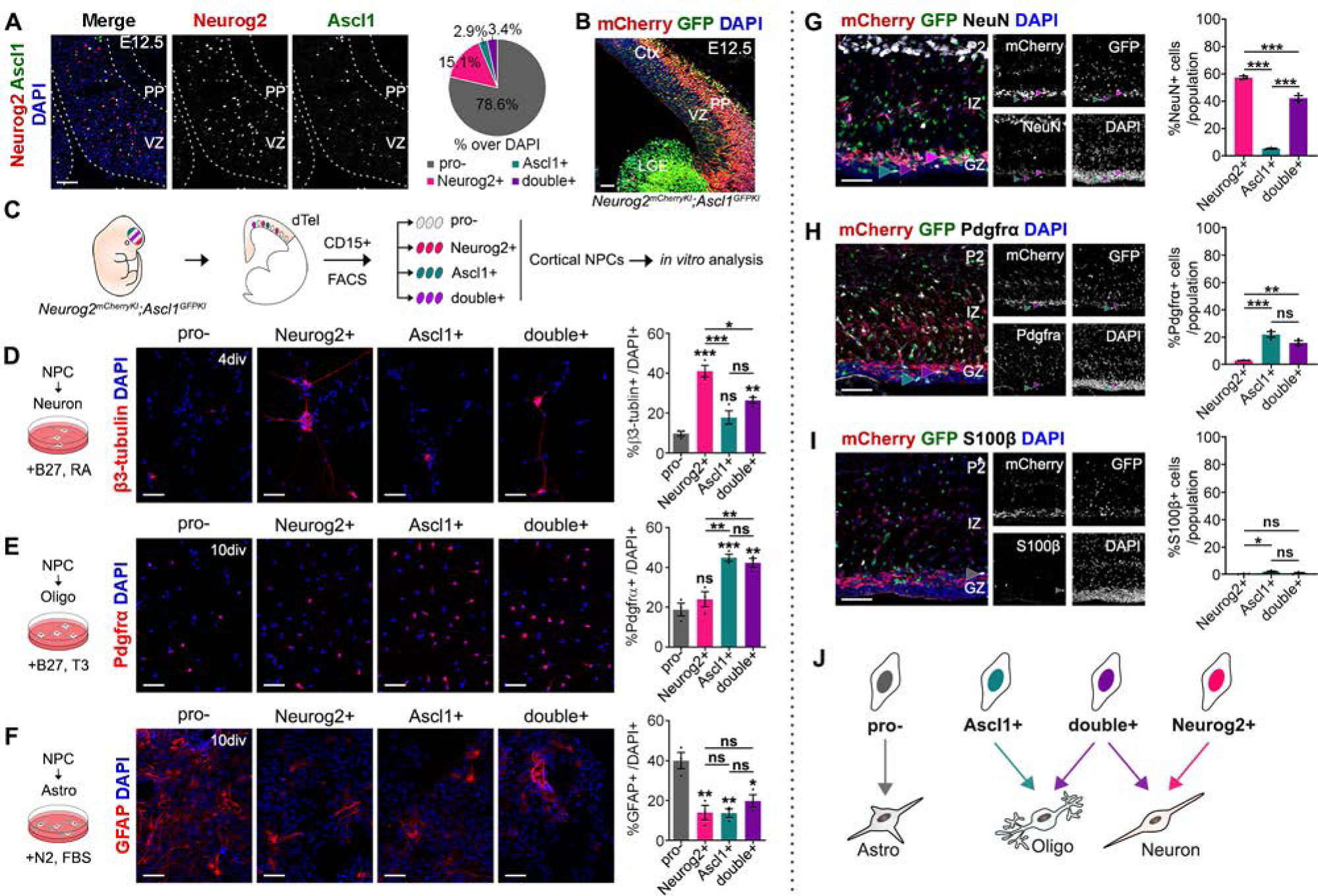
Neurog2 and Ascl1 expression defines four cortical NPC pools with distinct lineage biases. (A) Co-immunolabeling of E12.5 mouse cortex with Neurog2 (red) and Ascl1 (green). Quantification of the proportion of DAPI^+^ NPCs expressing Neurog2 and Ascl1 (N=3). (B) mCherry and GFP expression in E12.5 *Neurog2^mCherry^*^KI^;*Ascl1^GFP^*^KI^ (double KI) transgenics. (C) Schematic of FACS isolation protocol of CD15^+^ cortical NPCs. (D-F) Directed differentiation of E12.5 CD15^+^ pro^−^, Neurog2^+^, Ascl1^+^ and double^+^ NPCs into β3-tubulin^+^ neurons after 4 DIV (N=3) (D); Pdgfrα^+^ oligodendrocytes after 10 DIV (N=3) (E); and GFAP^+^ astrocytes after 10 DIV (N=3) (F). (G-I) Co-immunolabeling of mCherry^+^ and/or GFP^+^ cells in P2 double KI cortices with NeuN (G), Pdgfrα (H) and S100β (I). (N=3 for all markers). All comparisons made with one-way ANOVA and Tukey’s post-hoc tests. p-values: ns – not significant, <0.05 *, <0.01 **, <0.001 ***. Scale bars in A-B,D-F,G-I = 50 µm. Ctx, cortex; LGE, lateral ganglionic eminence; PP, preplate; VZ, ventricular zone. See also Figure S1.

To identify and study NPCs expressing the two proneural genes, we generated *Neurog2^mCherryKI^* knock-in (KI) mice (Figure S1A-D) that were crossed with *Ascl1^GFPKI^* transgenics (Leung et al., 2007), creating double KIs. In E12.5 double KIs, mCherry and GFP were expressed in the VZ, where NPCs reside, and persisted in postmitotic neurons (Figure 1B), acting as a short-term lineage trace. Double KIs can thus identify pro^−^ (no reporter), Neurog2^+^ (mCherry^+^), Ascl1^+^ (GFP^+^) and double^+^ (mCherry^+^GFP^+^) lineages.

We used this short-term lineage tracing system to ask how *Neurog2* and *Ascl1* expression correlates with cell fate decision by cortical NPCs using *in vitro* differentiation assays. Cortical NPCs sequentially give rise to neurons (between embryonic day (E) 10 – E17 (Takahashi et al., 1999), astrocytes (beginning at E16) and oligodendrocytes (beginning early postnatally). To assess the developmental potential of the four NPC pools in the E12.5 neurogenic phase, FACS was used to isolate pro^−^, Neurog2^+^, Ascl1^+^, and double^+^ NPCs from double KI cortices using mCherry and GFP expression. We further selected for CD15 expression to enrich for NPCs over neurons (Ballas et al., 2005) (Figure 1C). CD15 marked a subset of CD133^+^ cells, another NPC marker; (Furutachi et al., 2015), but as CD15^+^ NPCs had more expression of VZ radial glia (RG) versus IPC markers, we used CD15 to sort NPCs (Figure S1E-K). Enriched expression of *Neurog2* and *Ascl1* was validated in E12.5 CD15^+^ pro^−^, Neurog2^+^, Ascl1^+^ and double^+^ NPCs by qPCR (Figure S1L).

After 4 days *in vitro* (DIV), sorted Neurog2^+^ NPCs gave rise to the most β3-tubulin^+^ neurons (Figure 1D), above baseline levels defined as neuronal number generated by the pro^−^ NPC pool. Conversely, Ascl1^+^ NPCs were biased towards generating Pdgfrα^+^ oligodendrocytes, above pro^−^ NPCs (Figure 1E). Interestingly, double^+^ NPCs produced neurons and oligodendrocytes above baseline levels suggesting that these cells are less lineage restricted (Figure 1D-E). Finally, GFAP^+^ astrocyte production was highest in pro^−^ NPCs (Figure 1F).

To confirm lineage biases *in vivo*, we directly examined double KI cortices at postnatal day (P) 2, when neurogenesis has ended and gliogenesis is in progress. Most *Neurog2-*lineage cells (mCherry^+^) co-labeled with NeuN, a pan-neuronal marker, which was also expressed in double^+^ lineage cells and to a much lesser extent in the *Ascl1-*lineage (GFP^+^) (Figure 1G). Instead, Pdgfrα co-labeled most cells in Ascl1^+^ and double^+^ lineages, while very few Neurog2^+^ lineage cells were Pdgfrα^+^ (Figure 1H). Finally, for all pro^+^ populations, few cells co-labeled with S100b, an astrocyte marker (Figure 1I).

Neurog2^+^ and Ascl1^+^ NPCs are thus biased towards neuronal and oligodendrocyte lineages, respectively, while double^+^ NPCs are less lineage restricted (Figure 1J).

### Neurog2 and Ascl1 form a cross-repressive toggle switch to control lineage commitment

Co-expression of Neurog2 and Ascl1 in less lineage-restricted NPCs was reminiscent of hematopoiesis, in which bipotent progenitors co-express pairs of cross inhibitory, lineage-specifying TFs and upon fate restriction convert to express a single dominant factor (Brand and Morrissey, 2020). To determine whether proneural double^+^ NPCs similarly convert to single^+^ NPCs over time, we performed time-lapse imaging on E14.5 double KI cortical slices over an ∼18 hr period (Figure 2A, Movie File S1). We observed that mCherry^+^GFP^+^ cells in the VZ preferentially converted to mCherry^+^ (55%) or GFP^+^ (33%) cells, with fewer remaining double^+^ (12%) (Figure 5A). Thus, a subset of double^+^ NPCs convert to Neurog2^+^ or Asc1l^+^ NPCs over time.

**Figure 2.**
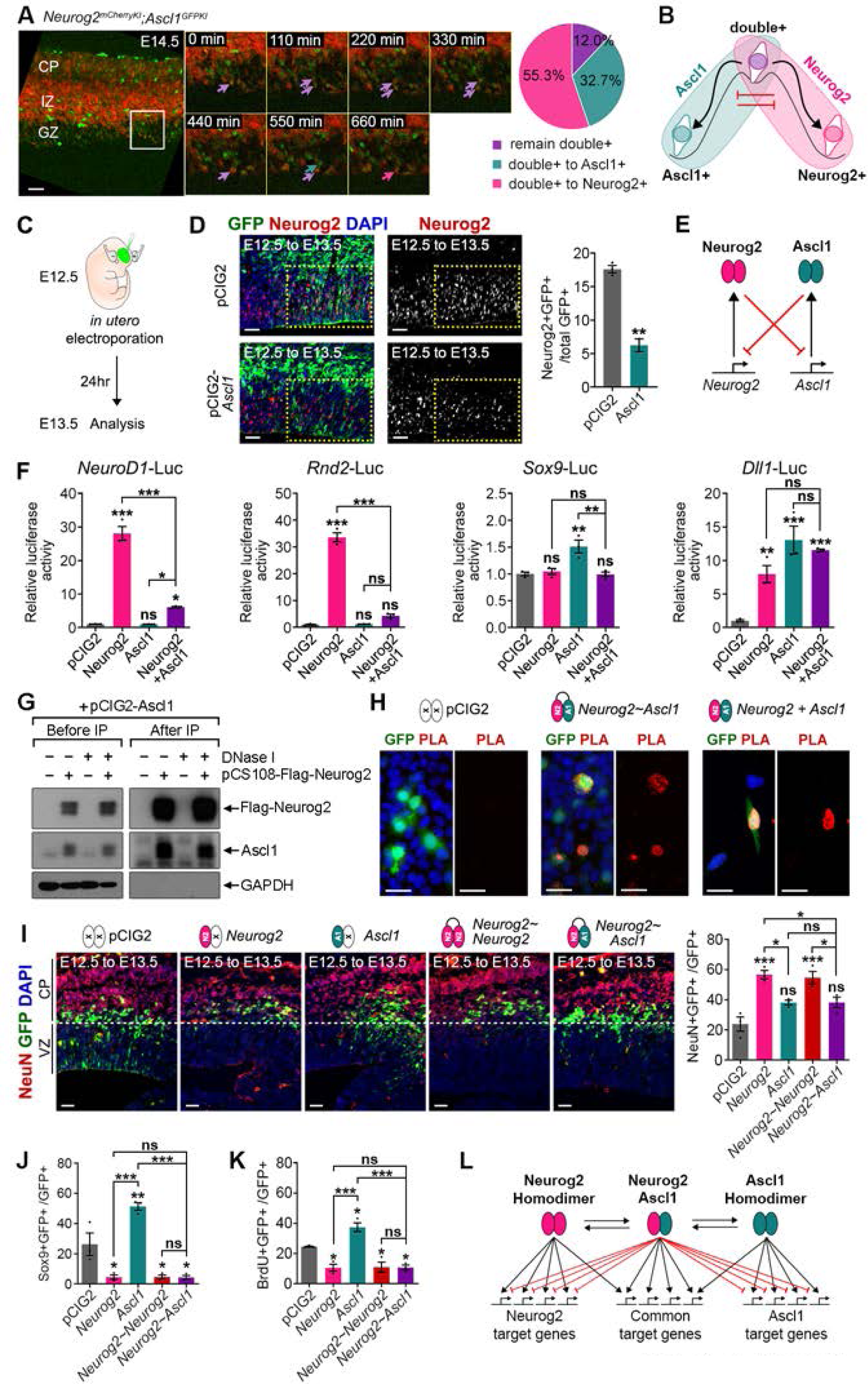
Neurog2 and Ascl1 are cross-repressive at the level of expression and function. (A) Time-lapse imaging of E14.5 *Neurog2^mCherryKI^;Ascl1^GFPKI^* cortical NPCs over 660 min (N=3). Arrows point to double^+^ NPCs (purple arrows) that become Ascl1^+^ (green arrow) or Neurog2^+^ (pink arrow). Quantitation of % double^+^ NPC conversion (N=3). (B) Proposed model of double^+^ lineage conversion to Neurog2^+^ and Ascl1^+^ NPCs. (C,D) Schematic of E12.5 to E13.5 electroporations (C). Electroporation of pCIG2 (GFP control) or *Ascl1*, followed by analysis and quantification of Neurog2^+^GFP^+^ cells (N=3 for pCIG2, N=2 for Ascl1) (D). Yellow boxes outline electroporated regions. (E) Neurog2 and Ascl1 cross-repress at the expression level. (F) Transcriptional reporter assays using *NeuroD1*, *Rnd2*, *Sox9*, and *Dll1* luciferase reporters (N=3 for each reporter and condition). (G) Co-immunoprecipitation of Ascl1 with FLAG-Neurog2 with or without DNaseI treatment *in vitro*. (H) Proximity ligation assay (PLA) on P19 cells transfected with pCIG2 (negative control), Neurog2∼Ascl1 (tethered) or Neurog2∼Ascl1 (tethered) using Ascl1 and Neurog2 antibodies. (I-K) E12.5 to E13.5 *in utero* electroporation of pCIG2, *Neurog2*, *Ascl1*, *Neurog2∼Neurog2* (tethered) or *Neurog2∼Ascl1* (tethered). Quantification of electroporated GFP^+^ cells that co-express NeuN (I), Sox9 (J), or incorporate BrdU after 30 min (K) (N=3 for all markers). (L) Schematic of Neurog2 and Ascl1 dimerization and functional cross-repression. Two-tailed Student’s t-test was used in C. One-way ANOVA with Tukey’s post-hoc tests were used for multiple comparisons in F,I-K. p-values: ns – not significant, <0.05 *, <0.01 **, <0.001 ***. Scale bars in A = 50 µm and D,H,I = 25 µm. CP, cortical plate; GZ, germinal zone; IZ, intermediate zone; VZ, ventricular zone. See also Figure S2.

In hematopoiesis, TF pairs are cross-repressive at the level of expression and function, prompting us to ask whether *Neurog2* and *Ascl1* function in a similar way (Figure 2B). In cortical NPCs, *Neurog2* is required and sufficient to repress *Ascl1* expression, which is upregulated in *Neurog2*^−/-^ brains (Fode et al., 2000). To examine whether conversely, *Ascl1* over-expression inhibits expression of *Neurog2*, E12.5 cortices were electroporated with pCIG2-*Ascl1,* revealing a reciprocal repression of *Neurog2* expression by *Ascl1* (Figure 2C-E).

We next questioned whether *Neurog2-Ascl1* also cross-repress at the level of function. We found that indeed *Neurog2* and *Ascl1* inhibit each other’s abilities to transactivate downstream target genes using transcriptional reporter assays with *Neurog2* (*NeuroD1*, *Rnd2*) and *Ascl1* (*Sox9*) target genes (Li et al., 2014; Li et al., 2012) (Figure 2F). In contrast, a *Dll1* reporter that is a target of both *Neurog2* and *Ascl1* was not repressed by co-expression (Figure 2F). Proneural TFs dimerize, suggesting that cross-repression could be mediated by Neurog2-Ascl1 heterodimer formation. We confirmed that Neurog2 and Ascl1 physically interact using immunoprecipitation (Figure 2G) and a proximity ligation assay (Figure 2H). To examine the *in vivo* consequence of *Neurog2-Ascl1* interaction, E12.5 cortices were electroporated with expression constructs for *Neurog2*, *Ascl1*, and tethered *Neurog2∼Neurog2* and *Neurog2∼Ascl1* heterodimers (Figure 2I). Both *Neurog2* and *Neurog2∼Neurog2* promoted neurogenesis above pCIG2 baseline levels 24 hr post-electroporation, while *Neurog2∼Ascl1* produced 1.5-fold fewer NeuN^+^GFP^+^ neurons relative to *Neurog2* or *Neurog2∼Neurog2,* indicative of an inhibitory effect of *Ascl1* on *Neurog2* function (Figure 2I). Similarly, *Neurog2* inhibited the ability of *Ascl1* to induce the expression of Sox9 (Figure 2J, S2A), an *Ascl1*-target gene and proliferative glioblast marker, and the uptake of BrdU (Figure 2K, S2B), which *Ascl1* induces (Li et al., 2014).

*Neurog2* and *Ascl1* are thus cross-repressive at the level of expression and function, providing mechanistic insight into why double^+^ NPCs are less lineage committed (Figure 2L).

### Proneural^−^, Neurog2^+^, Ascl1^+^, and double^+^ cortical NPCs have unique transcriptomes

Our goal was to determine how proneural genes control neurogenic regularity in lissencephaly, but our finding that their expression defines four NPC pools with distinct differentiation properties suggested that each NPC pool is likely under different regulatory control. To understand the molecular basis for the unique cellular properties of pro^−^, Neurog2^+^, Ascl1^+^ and double^+^ NPCs, we performed RNA-seq analyses on E12.5 CD15^+^ FACS-purified NPCs. Principle component analysis (PCA) revealed pro^−^ and Ascl1^+^ NPCs were the most divergent, while Neurog2^+^ and double^+^ NPCs were the most similar, albeit still distinguishable (Figure 3A). Dissimilarity of the four NPC pools was confirmed using PCA analyses of transcript counts for 114 stem cell genes from a custom Nanostring codeset (Figure S3A, Table S1). Examination of a small subset of markers for cortical RG (*Sox2, Nestin, Pax6, BLBP, GLAST*), IPCs (*Tbr2*), glutamatergic (*Tbr1, Vglut1, Vglut2, Tubb3*) and GABAergic (*Gad1, Gad2*) neurons, oligodendrocytes (*Olig1, Olig2*) and astrocytes (*S100b, Aldh1l1, GFAP*) also stratified the four pools; pro^−^ and Ascl1^+^ NPCs were enriched in RG markers (Figure 3B, S3B), while Neurog2^+^ and double^+^ NPCs expressed higher levels of glutamatergic neuronal and IPC markers (Figure 3B, S3C-E). Notably, Ascl1^+^ NPCs expressed only very low levels of GABAergic and oligodendrocyte markers (Figure S3F-G), while astrocyte markers were not expressed in any NPCs at E12.5 (RPKM<1 in all population, data not shown). These gene expression profiles are consistent with the increased propensity of Neurog2^+^ and double^+^ NPCs to differentiate into neurons, while pro^−^ and Ascl1^+^ NPCs retain a RG identity, with low transcript counts for differentiation genes.

**Figure 3.**
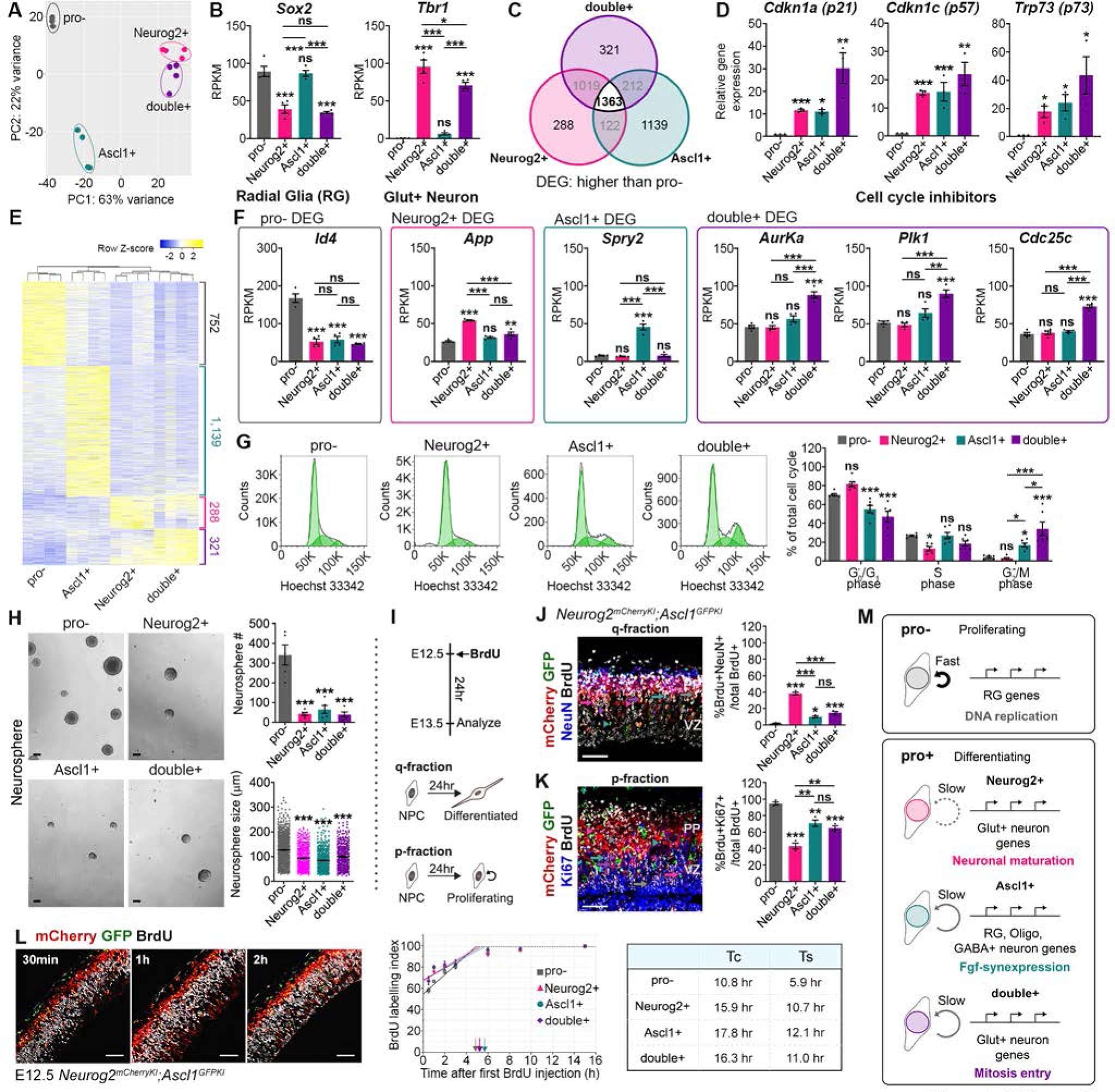
Pro^−^, Neurog2^+^, Ascl1^+^, and double^+^ cortical NPCs have unique transcriptomes and cell cycle properties. (A) PCA analysis of RNA-seq data from E12.5 CD15^+^ pro^−^, Neurog2^+^, Ascl1^+^ and double^+^ NPCs (N=4). False discovery rate (FDR) less than 5%. (B) RPKM values of select RG and neuronal markers. (C) Venn diagram of DEGs >1.5-fold higher in in pro^+^ versus pro^−^ NPCs (p-value <0.05). (D) qPCR validation of negative cell cycle regulators *Cdkn1a*, *Cdkn1c* and *Trp73* in E12.5 CD15^+^ NPCs (N=3), normalizing pro^−^ NPCs to 1. (E) Heatmap showing DEGs in E12.5 CD15^+^ pro^−^, Neurog2^+^, Ascl1^+^ and double^+^ NPCs. (F) RPKM values of select genes enriched in pro^−^, Neurog2^+^, Ascl1^+^ and double^+^ E12.5 CD15^+^ NPCs. (G) Cell cycle analysis of E12.5 CD15^+^ pro^−^, Neurog2^+^, Ascl1^+^ and double^+^ NPCs stained with Hoechst 33342. Quantitation of the proportions of NPCs in G_0_/G_1_, S and G_2_/M phases (N=6) (H) Neurospheres derived from E12.5 CD15^+^ pro^−^, Neurog2^+^, Ascl1^+^ and double^+^ NPCs and imaged and quantitated (number, size) after 7 DIV. (N=5). (I-K) Schematic of q- and p-fraction analysis (I). Co-labeling E12.5 double KI cortices with mCherry, GFP, NeuN and BrdU to calculate q-fraction (%BrdU^+^NeuN^+^/ BrdU^+^ cells) (N=3) (J). Co-labeling of mCherry, GFP, Ki67 and BrdU to calculate p-fraction (%BrdU^+^Ki67^+^/ BrdU^+^ cells) (N=3) (K). (L) Cumulative BrdU labeling to calculate cell cycle time (Tc) and S-phase length (Ts) in E12.5 double KIs. Co-labeling of mCherry, GFP and BrdU after 0.5, 1, and 2 hr of BrdU exposure (N=3 each). Plot of BrdU labelling index (% BrdU^+^ nuclei) and table summarizing calculated Tc and Ts values. Labeling index saturation points indicated by color-coded arrows. (M) Summary of proliferation properties and lineage biases. Two-tailed Student’s t-tests were used in D. One-way ANOVA with Tukey’s post-hoc tests in B,F,H,J,K and two-way ANOVA with Tukey’s post-hoc test in G were used for multiple comparisons. p-values: ns – not significant, <0.05 *, <0.01 **, <0.001 ***. Scale bar in H = 100 µm and J-L = 50 µm. Glut^+^, glutamatergic; GABA^+^, GABAergic; Oligo, Oligodendrocyte; PP, preplate; VZ, ventricular zone. See also Figure S3.

We next searched for differentially expressed genes (DEGs) signatory of the pro^+^ NPC phenotype, encompassing Neurog2^+^, Ascl1^+^ and double^+^ NPCs (Figure 3C, Table S2). Of the 1363 genes enriched in pro^+^ versus pro^−^ NPCs, several closely clustered negative regulators of the cell cycle were upregulated in the pro^+^ cohort (*Cdkn1a*, *Cdkn1c*, *Trp73*) (Figure S3H), confirmed by qPCR (Figure 3D). Accordingly, overexpression of *Cdkn1a*, *Cdkn1c* and *Trp73* in E12.5 NPCs *in vitro* inhibited the formation of primary neurospheres (Figure S3I-K).

To better understand NPC differences, we also examined uniquely expressed genes in each NPC pool (Figure 3E, Table S2). The 752 DEGs in pro^−^ NPCs were enriched in GO terms related to DNA replication and negative regulation of neurogenesis (e.g. *Mcm3*/5, *Rpa1*, *Id4*, Figure 3F, S3L), consistent with their higher proliferative potential. The 288 DEGs specific to Neurog2^+^ NPCs were enriched for GO terms related to neuronal maturation (e.g. synaptic potential, *App*, *Tmod2;* Figure 3F, S3M), while the 1139 Ascl1^+^ specific DEGs were enriched in GO terms related to various signaling pathways, such as MAPK (e.g. *Fgf15/18*, *Spry1/2*; Figure 3F, S3N), which when activated converts *Ascl1* to a pro-gliogenic determinant (Li et al., 2014). Finally, of the 321 DEGs enriched in double^+^ NPCs, GO terms related to mitosis entry were highly enriched (e.g. *Aurka*, *Plk1*, *Cdc25c;* Figure 3F, S3O). Consistent with this finding, cell cycle profiling using flow cytometric measures of DNA content revealed a higher proportion of double^+^ NPCs in G_2_/M phase, at the expense of cells in G1 (Figure 3G).

*Neurog2* and *Ascl1* expression thus defines four NPC pools with distinct transcriptomes, which were further suggestive of cell cycle differences.

### Proneural^+^ NPCs have longer cell cycles and an enhanced propensity to differentiate

In lissencephalic rodent brains, neurogenic continuity is the summation of coordinated decisions by individual cortical NPCs to differentiate or proliferate. Transcriptomic analyses suggested there may be differences in proliferation potential amongst cortical NPCs expressing *Neurog2* and/or *Ascl1*, which we first assessed with a neurosphere assay. Sorted CD15^+^ pro^−^, Neurog2^+^, Ascl1^+^ and double^+^ NPCs were plated at clonal density, and after 7 days *in vitro*, neurosphere number and size were enumerated. Pro^−^ NPCs formed an order of magnitude more neurospheres that were larger in size than any of the pro^+^ populations (Figure 3H), suggesting that pro^−^ NPCs includes the most sphere-forming NSCs, and that these cells and/or their progeny have an enhanced proliferative capacity. Conversely, all three pro^+^ NPCs gave rise to fewer and smaller neurospheres (Figure 3H), negatively correlating proneural gene expression with stemness.

To further examine the association between proneural gene expression and NPC choice to proliferate or differentiate, we calculated proliferative (p) and differentiative (leaving or q) fractions for the four cortical NPC pools. BrdU was administered to pregnant double KIs at E12.5 and embryos were sacrificed 24 hrs later (Figure 3I). The q-fraction (%BrdU^+^NeuN^+^/BrdU^+^ cells) of the pro^−^ pool was negligible, suggesting these NPCs have a low probability to differentiate, while >30% of all pro^+^ NPCs exited the cell cycle, with Neurog2^+^ NPCs the most likely to differentiate into NeuN^+^ neurons (Figure 3J). Conversely, p-fractions were highest in pro^−^ NPCs (%BrdU^+^Ki67^+^/BrdU^+^ cells), confirming a bias to stay in the cell cycle, while p-fractions were lower for all pro^+^ NPCs, especially the Neurog2^+^ NPC pool (Figure 3K). Proneural TF expression, especially Neurog2, is thus a strong correlate of cell cycle exit and neurogenesis.

Finally, we asked whether the increased incidence of cell cycle exit by pro^+^ NPCs correlated with an increase in cell cycle length, which is generally associated with differentiation (Dalton, 2015). To measure total cell cycle (Tc) and S-phase (Ts) lengths, we used cumulative BrdU labeling, administering BrdU every three-hours to E12.5 double KI animals (Figure 3L). Enumeration of BrdU+ cells revealed that all pro^+^ NPCs had a longer Tc and Ts compared to pro^−^ NPCs, but were not different from one another.

Thus, in aggregate, pro^+^ NPCs have longer cell cycle lengths that are conducive to differentiation, but even so, only a subset of these NPCs differentiates at any given time, with singular Neurog2 expression the best determinant of neuronal differentiation (Figure 3M).

### Pro^−^, Neurog2^+^, Ascl1^+^, and double^+^ NPCs display differences in chromatin accessibility

As NPCs differentiate, they undergo a continuum of semi-stable cell fate transitions that increasingly restrict lineage potential (Telley et al., 2019). Fate restriction is accompanied by the closure of chromatin sites associated with multipotency and the opening of sites where lineage-specifying TFs can bind, culminating in a stable epigenome signatory of a terminally differentiated cell fate (Stergachis et al., 2013) (Figure 4A). TFs are often architects of epigenetic change as they can recruit chromatin modifiers to target sites (Brand and Morrissey, 2020). We thus performed ATAC-seq on E12.5 CD15^+^ pro^−^, Neurog2^+^, Ascl1^+^ and double^+^ NPCs to compare open chromatin associated with proneural gene expression (Figure 4A’). A genome-wide comparison of ‘open’ ATAC-seq peaks at transcription start sites (TSS) revealed that they were most abundant in double^+^ NPCs, followed by Ascl1^+^, Neurog2^+^ and pro^−^ NPCs (Figure 4B). Proneural TF expression is thus associated with increased chromatin accessibility at TSSs, especially when Neurog2 and Ascl1 are both expressed, consistent with the enhanced lineage potential of double^+^ NPCs.

**Figure 4.**
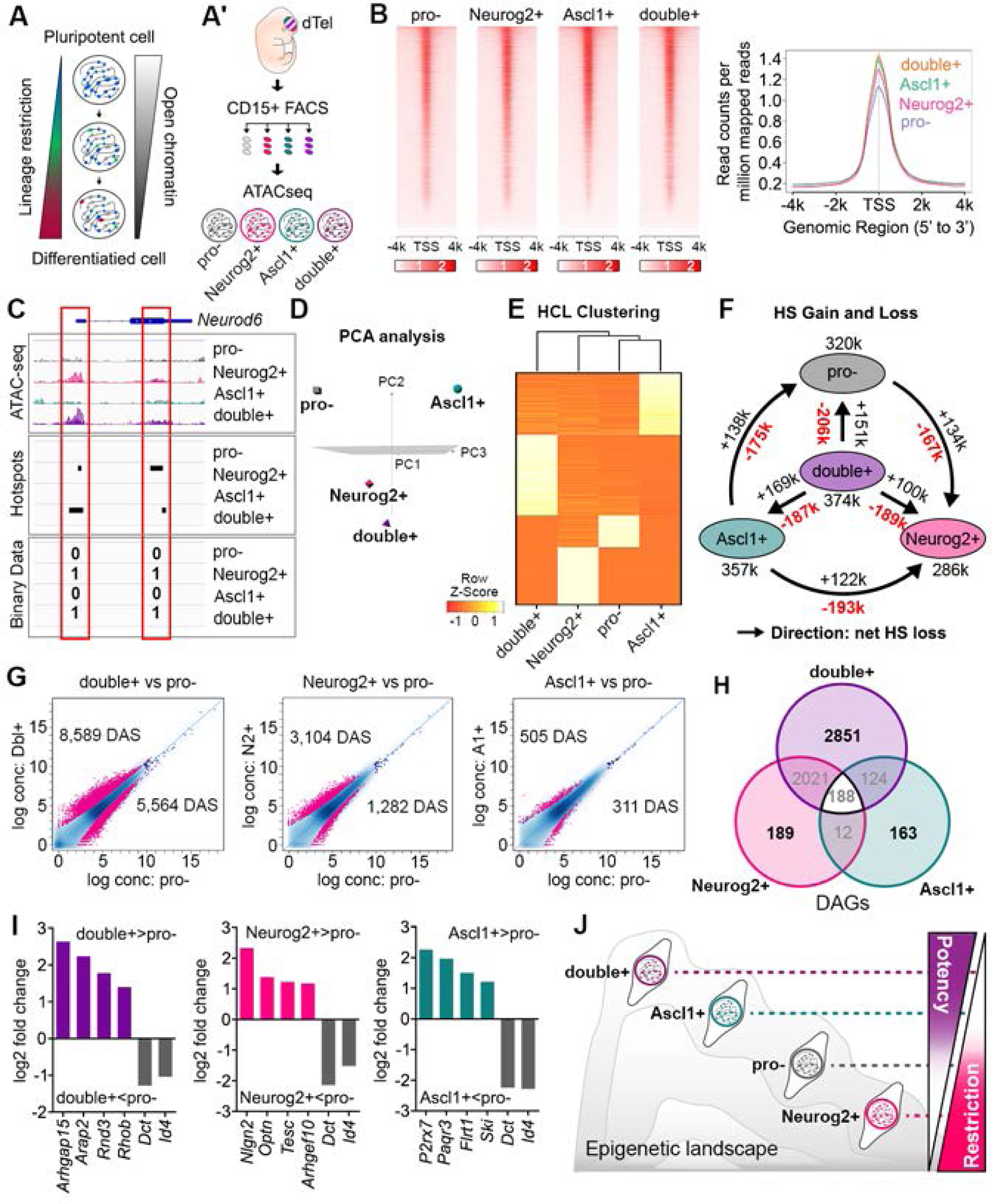
Pro^−^, Neurog2^+^, Ascl1^+^, and double^+^ cortical NPCs have distinct epigenetic landscapes. (A,A’) Schematic of gradual acquisition of lineage biases coincident with a global loss of open chromatin (A). Experimental paradigm for ATAC-seq analysis of open chromatin in E12.5 CD15^+^ pro^−^, Neurog2^+^, Ascl1^+^ and double^+^ NPCs (N=3) (A’). (B) Heatmap of ATAC-seq TSS peaks in each NPC pool. Read counts per million mapped reads near TSS. (C-F) Hotspot (HS) analysis used to quantitate ATAC-seq data, showing sample analysis on *Neurod6* locus (C). PCA of high confidence HSs present in all three biological replicates for each NPC pool (D). Hierarchical clustering of gene assigned HSs in each NPC pool (E). HS gain and loss in pairwise comparisons (F). Arrows show direction of net HS loss, suggestive of lineage restriction. (G-I) Diffbind analysis of ATAC-seq data showing differential accessible sites (DAS) in pairwise comparisons between pro^−^ and double^+^, Ascl1^+^ and Neurog2^+^ NPCs (G). Venn diagram showing the number of NPC pool-specific differentially accessible genes (DAG) compared to pro^−^ NPCs (H). Fold differences in open chromatin for select gene sets, showing comparisons between pro^−^ NPCs versus double^+^, Ascl1^+^ or Neurog2^+^ NPCs (I). (J) Epigenetic landscape model depicting degree of lineage restriction based on comparison of open chromatin sites. See also Figure S4.

**Figure 5.**
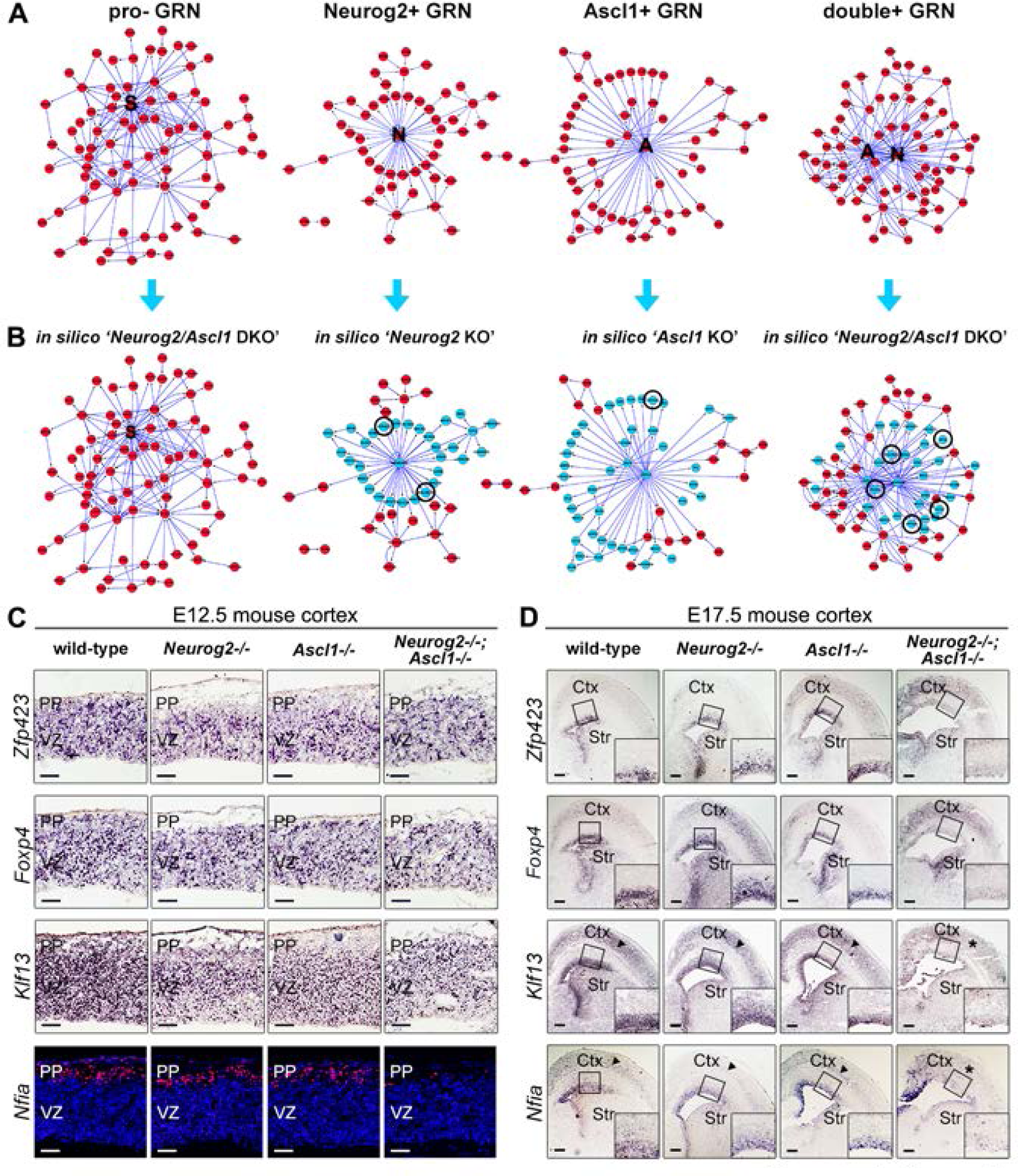
*In silico* and *in vivo* analyses of gene regulatory networks associated with pro^−^, Neurog2^+^, Ascl1^+^, and double^+^ cortical NPCs. (A) GRNs associated with E12.5 CD15^+^ pro^−^, Neurog2^+^, Ascl1^+^ and double^+^ NPCs, generated by combining RNA-seq, ATACseq and *in vivo* ChIP-seq data (Sessa et al., 2017). (B) *In silico* perturbations showing effects of a ‘*Neurog2/Ascl1* DKO’ on the pro^−^ GRN, ‘*Neurog2* KO’ on the Neurog2^+^ GRN, ‘*Ascl1* KO’ on the Ascl1^+^ GRN, and ‘*Neurog2* and ‘*Neurog2/Ascl1* DKO’ on the double^+^ GRN. Blue nodes depict a predicted loss of gene expression. Genes selected for further analysis are circled. (C,D) Immunofluorescence and RNA *in situ* hybridization analysis of the expression profiles of three predicted Neurog2/Ascl1 target genes, including *Zfp423*, *Foxp*4, *Klf13* and *Nfia* in E12.5 (C) and E17.5 (D) wild-type, *Neurog2*^−/-^, *Ascl1*^−/-^ and *Neurog2*^−/-^;*Ascl1*^−/-^ cortices. Insets are 2x magnifications of boxed regions in D. Scale bars in C = 50 µm and D = 200 µm. Ctx, neocortex; PP, preplate; Str, striatum; VZ, ventricular zone. See also Figure S5.

Chromatin is extensively reorganized during lineage restriction, including outside TSSs (Dixon et al., 2015). To interrogate genome wide differences in open chromatin sites, we employed a hotspot (HS) algorithm, assigning a value of “1” to open sites, designated HSs (John et al., 2011) (Figure 4C). PCA of total HSs revealed distinct DNA regulatory landscapes for all four NPC pools, with double^+^ and Neurog2^+^ NPCs most closely related, and Ascl1^+^ and pro^−^ NPCs the most divergent (Figure 4D), similar to transcriptomic comparisons (Figure 3A). Strikingly, hierarchical clustering analysis for nearest gene assigned HSs showed that each of the four NPC pools had distinct, almost non-overlapping sites of open chromatin, with double^+^ NPCs being the most divergent (Figure 4E). Furthermore, an analysis of all HSs, whether gene assigned or not, revealed the highest HS number in double^+^ NPCs (374K), followed by Ascl1^+^ NPCs (357K), as seen in total TSS peak comparisons (Figure 4F). However, pro^−^ NPCs (320K) had more HSs compared to Neurog2^+^ NPCs (286K), opposite to the TSS peaks, indicating that even though Neurog2^+^ NPCs have fewer sites of open chromatin, more of these sites are in TSSs (Figure 4F).

An increase in open chromatin sites correlates with more ‘active’ TF binding, and is also associated with multipotency, while conversely, reduced chromatin accessibility is associated with the transition to a more lineage restricted state (Klemm et al., 2019; Stergachis et al., 2013). We used ATAC-seq data to query lineage restriction by examining HS gain and loss in pairwise comparisons between each NPC pool, following the assumption that more HSs are lost than gained during lineage restriction (Stergachis et al., 2013). We found that double^+^ NPCs had a net loss of HSs in all three pairwise comparisons (Figure 4F). Conversely, in all pairwise comparisons, Neurog2^+^ NPCs never lost HSs, suggesting they are more fate restricted. Finally, pro^−^ (net loss in 2 comparisons) and Ascl1^+^ (net loss in one comparison) NPCs had intermediate phenotypes (Figure 4F).

HS analysis evaluates number of open chromatin sites, but does not provide information on the degree of accessibility of individual sites. We therefore applied a Diffbind analysis to identify differential accessible sites (DAS). In pairwise comparisons to pro^−^ NPCs, double^+^ NPC had the most DAS (FDR <0.10), followed by Neurog2^+^ and Ascl1^+^ NPCs (Figure 4G, Table S3). These data support the idea that Ascl1^+^ and pro^−^ NPCs have the most closely related open chromatin structure, while Neurog2^+^ and double^+^ NPCs are more distantly related. Finally, within the DAS, we identified differential accessible genes (DAG) unique to each NPC pool (Figure 4H); double^+^ NPCs had the most unique DAGs (2,851 total), which were enriched in actin filament assembly and GTPase signal transduction (e.g. *Arhgap15*, *Arap2*, *Rnd3*, *Rhob;* Figure 4I, S4A), while the remaining NPC pools had similar numbers of DAGs: Neurog2^+^ DAGs (189 total) in regulation of unfolded proteins and protein transport which are involved in neuroprotection (e.g. *Nlgn2*, *Optn*, *Tesc, Arhgef10;* Figure 4I, S4B), Ascl1l^+^ DAGs (163 total) were enriched in negative regulation of TGFβ and MAPK signaling (e.g. *P2rx7*, *Paqr3*, *Flrt1*, *Ski;* Figure 4I, S4C), and pro^−^ DAGs (64 total) in transcription and gene expression, including TF involved in NPC proliferation and stemness (e.g. *Dct*, *Id4;* Figure 4I, S4D-E).

Our data support a continuum of lineage restriction, from lowest to highest: double^+^< Ascl1^+^< pro^−^< Neurog2^+^ NPCs (Figure 4J), supporting the idea that double^+^ NPCs have the highest developmental potency and least lineage restriction.

### Distinct gene regulatory networks that define each NPC pool are deregulated in *Neurog2^−/−^;Ascl1^−/−^* cortices

Cell fate potential is defined by the repertoire of expressed TFs together with the accessibility of their cognate binding sites (i.e. chromatin opening) (Brand and Morrissey, 2020). By combining transcriptomic and epigenomic data, gene regulatory networks (GRNs) that maintain cell states can be visualized (Okawa et al., 2015). To visualize GRNs specific to pro^−^, Neurog2^+^, Ascl1^+^ and double^+^ NPCs, we combined differential expression data from RNA-seq and ATAC-seq datasets and incorporated published *in vivo* Neurog2 and Ascl1 ChIP-seq data (Sessa et al., 2017) (Figure 5A). Neurog2 and Ascl1 formed central hubs in their respective GRNs, and in the double^+^ GRN, whereas Sox2, an RG marker (Bayraktar et al., 2014), formed a central regulatory hub in pro^−^ GRNs (Figure 5A).

To understand the role of Neurog2 and Ascl1 in sustaining cell identities, we performed *in silico* perturbations to identify GRN hubs dependent on these factors. Strikingly, the double^+^ GRN lost the least number of total nodes (44.9%) in a simulated ‘*Neurog2/Ascl1* double KO (DKO)’, (Figure 5B), while single ‘*Neurog2* KO’ and ‘*Ascl1* KO’ simulations had even less of an effect on the double^+^ GRN (17.4% of total nodes lost by ‘*Neurog2* KO’, 8.7% of total nodes lost in by ‘*Ascl1 KO*’ (Figure S5A). Instead, single KO simulations were most disruptive on their corresponding GRNs; the Neurog2^+^ GRN lost 58.5% of total nodes in a ‘*Neurog2* KO’ and the Ascl1^+^ GRN lost 70.0% of total nodes in an ‘*Ascl1* KO’ (Figure 5B). Finally, as expected, the pro^−^ GRN was not affected by ‘*Neurog2* KO’ or ‘*Ascl1* KO’ simulations as these genes are not present (Figure 5B). These data support the idea that the four NPC pools are maintained by distinct GRNs and suggest that *Neurog2* and *Ascl1* largely compensate for each other to maintain the double^+^ GRN.

To validate the GRNs, we examined the expression of putative deregulated genes in E12.5 *Neurog2^−/-^* and *Ascl1^−/-^* single and double KOs. Most of the GRN genes analysed were expressed in the cortical VZ at E12.5 and were not noticeably perturbed in single or double KOs, including *Zfp423* (common to all GRNs), *FoxP4* (double^+^ GRN), and *Klf13* (double^+^ GRN), while GRN genes expressed in the cortical preplate, *Nfia* (double^+^ GRN) and *Bhlhe22* (Neurog2^+^ and double^+^ GRN), were both down-regulated only in *Neurog2^−/-^;Ascl1^−/-^* cortices (Figure 5C, S5B). Notably, by E17.5, a more striking phenotype was observed, with *Zfp423, FoxP4, Klf13* and *Nfia* all downregulated in *Neurog2^−/-^;Ascl1^−/-^* cortical progenitors (Figure 5D), correlating with a striking depletion of Pax6^+^ and Sox9^+^ RG and Tbr2^+^ IPCs in these DKO brains (Figure S5C-E). These data suggest that Neurog2 and Ascl1 largely compensate for one another to regulate cortical gene expression and to maintain the NPC pool, further validating the importance of their combined roles.

### Double^+^ NPCs are required for neurogenic symmetry and a lissencephalic phenotype

The unique features of double^+^ NPCs raised the question of whether they have an important role *in vivo*, which we addressed by developing a split-Cre lineage tracing system (Hirrlinger et al., 2009). N- and C-termini of Cre were knocked into *Neurog2* (*Neurog2^NCre^*) and *Ascl1* (*Ascl1^CCre^*) loci, respectively, allowing reconstitution of full-length, functional Cre only in double^+^ NPCs (Figure 6A). To trace the progeny of double^+^ NPCs, *Neurog2^NCre^*;*Ascl1^CCre^*^KI^ (hereafter split-Cre) mice were crossed with *Rosa-zsGreen* or Rosa-*tdTomato* reporters, creating ‘triple transgenics’. Recombined tdTomato^+^ cells were first detectable in the neuronal preplate at E12.5 and scattered throughout the cortex in small numbers at P0 (Figure S6A). At P7, zsGreen^+^ cells co-labeled with both NeuN (neurons) and Pdgfrα (oligodendrocytes) (Figure 6B-C), confirming the bipotency of double^+^ NPCs observed in short-term lineage tracing (Figure 1).

**Figure 6.**
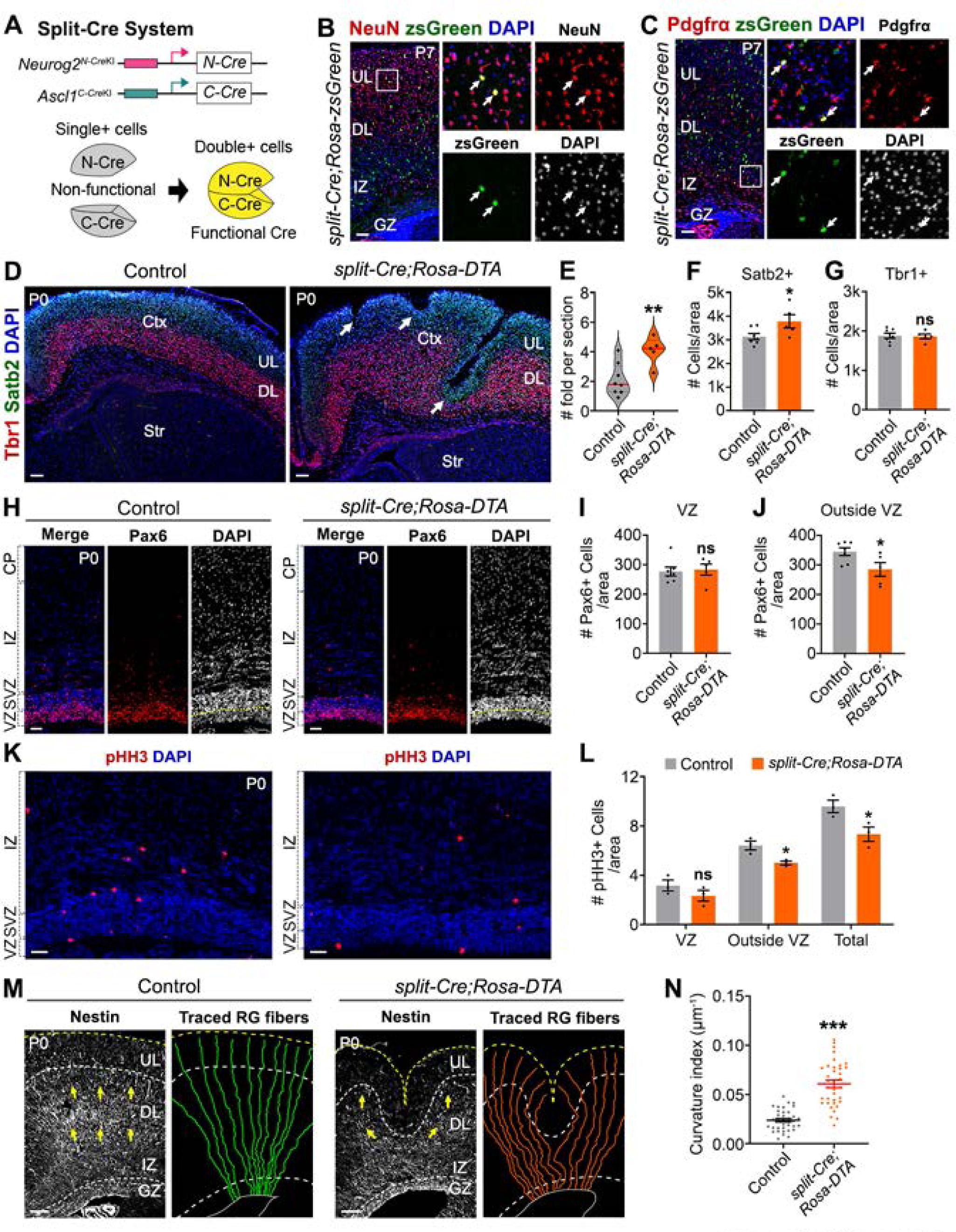
Double^+^ NPCs are essential to sustain regular neurogenesis and prevent cortical folding. (A) Knock-in (KI) strategy to generate *Neurog2/Ascl1* split-Cre transgenics. (B,C) Lineage tracing in P7 split-Cre;Rosa-zsGreen cortices, showing co-staining of zsGreen with NeuN (B) and Pdgfrα (C). Insets are 4x magnification of boxed areas. Arrows mark double^+^ cells. (D-G) P0 control and split-Cre;Rosa-DTA ‘deletor’ mice cortices immunostained with Tbr1 and Satb2 (D). Arrows in D mark cortical folds. Quantification of folds per section (N=8 control; N=5 split-cre;Rosa-DTA animals) (E). Quantification of Satb2^+^ (F) and Tbr1^+^ (G) neurons (N=7 control; N=5 split-cre;Rosa-DTA). (H-J) Immunostaining and quantification of Pax6^+^ NPCs in (I) and outside (J) the VZ in P0 control and split-Cre;Rosa-DTA cortices (N=7 control; N=5 split-Cre;Rosa-DTA). (K-L) Immunostaining (K) and quantification (L) of pHH3^+^ NPCs in and outside the VZ, and total in P0 control and split-Cre;Rosa-DTA cortices (N=3 each genotype). (M,N) Nestin immunostaining of P0 control and split-Cre;Rosa-DTA cortices (M). Arrows mark direction of RG projections. Representative aRG fibers are traced. Quantification of curvature index of individual RG (N=3, n=12 for each genotype) (N). Two-tailed Student’s t-tests were used in E-G, I-J, L and N. p-values: ns – not significant, <0.05 *, <0.01 **, <0.001 ***. Scale bars in B-D,M = 100 µm, and in H,K = 50 µm. Ctx, neocortex; DL, deep layer; GZ, germinal zone; IZ, intermediate zone; P, postnatal day; Str, striatum; SVZ, subventricular zone; UL, upper layer; VZ, ventricular zone. See also Figure S6.

To assess the requirement for double^+^ NPCs, we deleted these cells by crossing split-Cre mice with *Rosa-DTA* transgenics (hereafter split-cre;Rosa-DTA), resulting in Cre-dependent expression of diptheria toxin subunit A (DTA) only in double^+^ NPCs. As expected, dying TUNEL^+^ cells were detected in the germinal zone of P0 split-cre;Rosa-DTA mice (Figure S6B). Moreover, deep cortical folds restricted to the neuronal layers were observed in P0 split-cre;Rosa-DTA cortices (Figure 6D-E). The formation of these folds was associated with an increased number of Satb2^+^ upper layer neurons at both E15.5 (Figure S6C-D) and P0 (Figure 6F), while the number of Tbr1^+^ deep-layer neurons was similar to controls at both stages (Figure S6E; 6G).

Expansion of upper layer neurons is a feature of gyrencephalic cortices and is associated with the appearance of oRG, a basal NPC pool present in only small number in murine cortices (Wang et al., 2011). To assess differences in apical and basal NPCs in P0 split-cre;Rosa-DTA cortices, we examined the expression of Pax6 (marks RG and transient in IPCs), Sox9 (marks RG) and Tbr2 (marks IPCs). While the total number of Pax6^+^ and Sox9^+^ RG and Tbr2^+^ IPCs did not change in P0 split-cre;Rosa-DTA cortices (Figure 6H-I, S6F-K), there was a decrease in Pax6^+^ NPCs outside the VZ (Figure 6J), correlating with a reduction in mitotic phospho-histone H3 (pHH3)^+^ basal NPCs (Figure 6K-L). Therefore, cortical folding in P0 split-cre;Rosa-DTA mice is not due to oRG expansion, and instead develops despite a reduction in basal NPCs.

RG fibers curve in the vicinity of gyri in folded cortices due to the asymmetric insertion of oRG fibers amongst the evenly distributed ventricular RG fibers (Llinares-Benadero and Borrell, 2019). Using Nestin to label RG, we evaluated the trajectory of RG fibers in P0 split-cre;Rosa-DTA mice, observing an increased curvature at the location of cortical folding (Figure 6M,N).

Altogether, deletion of double^+^ NPCs results in the production of more upper layer neurons and cortical folding that is associated with a curvature of the radial glial scaffold that is not due to oRG expansion.

### Double^+^ NPCs regulate Notch signaling

A possible reason for the discontinuity of RG fibers in P0 split-cre;DTA cortices was a change in the pattern of NPC differentiation, as NPCs retract their RG fiber during differentiation. Proneural TFs cell autonomously induce NPC differentiation and non cell autonomously induce neighboring NPCs to proliferate via Notch signaling (Wilkinson et al., 2013). Proneural TFs regulate Notch signaling by transactivating the Notch ligands, *Dll1* and *Dll3* (Castro et al., 2006). To determine whether the pattern of Notch signaling could be regulated by *Neurog2* and *Ascl1*, we assessed Notch activity in the four NPC pools. Strikingly, from our RNA-seq data, *Dll1* and *Dll3* were expressed at the highest levels in Neurog2^+^ and double^+^ NPCs, while the *Notch2* receptor and two downstream effectors, *Hes1* and the Notch intracellular domain (NICD), were all elevated in pro^−^ and Ascl1^+^ NPCs (Figure 7A,B). Double^+^ and Neurog2^+^ NPCs are thus a rich source of Notch ligands, while Ascl1^+^ and pro^−^ NPCs respond to Notch signaling.

**Figure 7.**
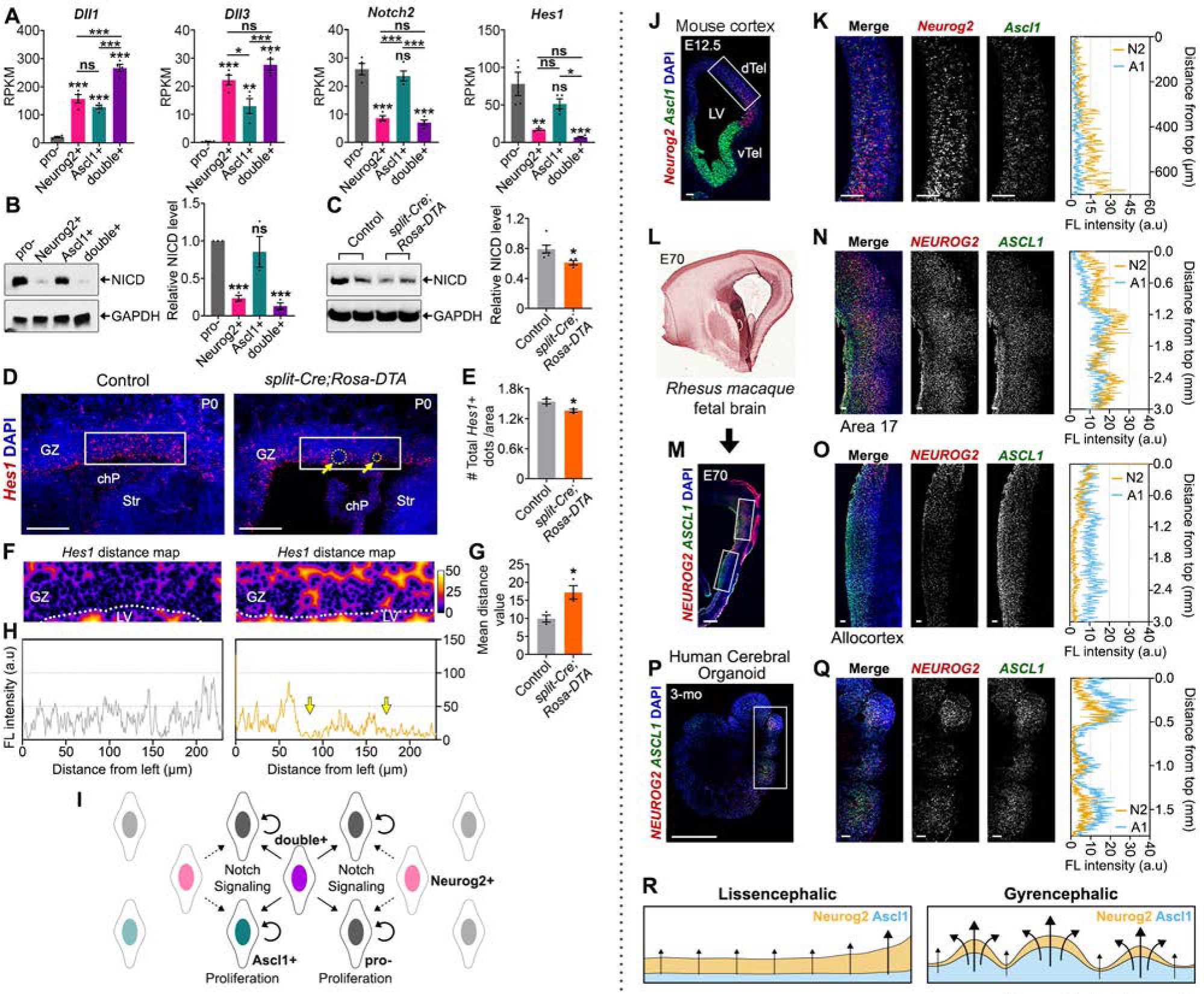
Double^+^ NPCs are Notch ligand expressing niche cells that are distributed evenly in mouse cortices and modular in gyrencephalic species. (A) RPKM values from RNAseq data for *Dll1*, *Dll3*, *Notch2* and *Hes1* in pro^−^, Neurog2^+^, Ascl1^+^ and double^+^ NPCs (N=4). (B) Western blots and densitometry of Notch intracellular domain (NICD) levels in sorted E12.5 CD15^+^ pro^−^, Neurog2^+^, Ascl1^+^ and double^+^ cortical NPCs. Densitometry normalized to GAPDH and to pro^−^ NPCs (set at 1) (N=3). (C) Western blots and densitometry of NICD levels in P0 control and split-Cre;Rosa-DTA cortices. Densitometry normalized to GAPDH (N=6 control; N=4 split-Cre;Rosa-DTA). (D-H) RNAscope analysis of *Hes1* transcript distribution in P0 control and split-Cre;Rosa-DTA cortices (D). Gaps in *Hes1* signals indicated by yellow arrows in D. Quantification of the number of *Hes1* dots (N=3 both genotypes) (E). Distance maps of minimum distance to the nearest *Hes1* transcript in the GZ in control and split-Cre;Rosa-DTA cortices (F). Quantification of mean distance (N=3) (G). Line plots show fluorescence intensity of *Hes1* signals along the x axis (H). (I) Model demonstrating double^+^ NPCs act as Notch ligand producing ‘niche’ cells that control NPC proliferation vs. neurogenesis. (J-Q) RNAscope analyses of *Neurog2* and *Ascl1* transcript distribution, first in E12.5 mouse cortex (J) and 3x magnification of boxed area (K). Parasagittal section of E70 macaque cortex. RNAscope analysis of *NEUROG2* and *ASCL1* of P70 macaque cortex (M), showing 4x magnification of boxed area 17 visual cortex (N) and allocortex (O). RNAscope analysis of *NEUROG2* and *ASCL1* in 3-month cerebral organoids (P) and 2x magnification of boxed area (Q). Line plots in K,N,O,Q shows quantification of fluorescence intensity. (R) Model of Neurog2/Ascl1 influence on cortical folding. One-way ANOVA with Tukey’s post-hoc tests were used for multiple comparisons in A. Two-tailed Student’s t-tests were used in B-C,E,G. p-values: ns – not significant, <0.05 *, <0.01 **, <0.001 ***. Scale bars in D,J-K,N,O,Q = 100 µm, and in M,P = 1 mm. chP, choroid plexus: dTel, dorsal telencephalon; GZ, germinal zone; LV, lateral ventricle: Str, striatum; vTel, ventral telencephalon.

The heterogeneity of Notch signaling in the four NPC pools suggested that removal of a critical Notch ligand source (i.e. double^+^ NPCs) would disrupt signaling. Indeed, overall NICD levels were reduced in P0 split-cre;DTA cortices (Figure 7C). Given that double^+^ NPCs are sparsely located throughout the VZ (Figure 1A), we reasoned that their removal could disrupt the spatial pattern of Notch signalling, which we assessed by examining the geometry of *Hes1* transcript distribution using RNAscope in situ hybridization. While there was a slight reduction in total *Hes1* RNAscope signals in P0 DTA triple transgenic cortices (Figure 7D,E), more telling was the increased spatial distance between *Hes1* dots, indicative of a sporadic disruption of Notch signaling centers (Figure 7F-H).

Double^+^ NPCs thus act as essential Notch-ligand expressing niche cells that are required to sustain a smooth pattern of neurogenesis and prevent cortical folding (Figure 7I).

### Proneural^+^ NPCs are distributed evenly in mouse and modular in gyrencephalic cortices

If proneural genes regulate cortical folding, we reasoned their expression patterns should differ in lissencephalic (rodent) versus gyrencephalic (primate) cortices. To test this assumption, we used RNAscope ISH to compare periodicity and relative expression levels. In E12.5 murine cortices, *Neurog2* and *Ascl1* transcripts were distributed evenly across the VZ, albeit with higher overall levels in lateral regions (Figure 7J,K), which are developmentally more mature (Caviness et al., 2009). In area 17 of the E70 macaque visual cortex, midway during neurogenesis (Rakic, 1974), the germinal zone was histologically ‘smooth’, whereas *NEUROG2* and *ASCL1* transcripts had a modular distribution (Figure 7L-N). In contrast, in the primate allocortex, where folding does not occur, *NEUROG2, ASCL1* transcripts were continuously distributed (Figure 7O). Notably, *PAX6*, which is also expressed in the cortical VZ, was evenly distributed in the visual and allo-cortices (Figure S7O-P), suggesting that the modular distribution of proneural gene transcripts was a unique feature.

As a surrogate measure of *NEUROG2* and *ASCL1* transcript distribution in human cortices, we also generated cerebral organoids from human embryonic stem cells (ESCs) (Lancaster and Knoblich, 2014). Cerebral organoids are pseudo-folded, as both the germinal zone and neuronal layers are folded (Qian et al., 2019). Nevertheless, in day 90 organoids, *NEUROG2* and *ASCL1* transcript distribution was modular, with higher expression correlating with sites of ‘pseudo-folds’ (Figure 7P,Q).

An additional distinction observed was that the *Neurog2:Ascl1* transcript ratio was ‘balanced’ in primate neocortex (0.92) and human cerebral organoids (0.81), which are folded and pseudo-folded, respectively, versus the dominant expression of one proneural gene in the lissencephalic rodent brain (1.84; *Neurog2-*rich) and primate allocortices (0.29; *ASCL1*-rich). In sum, *Neurog2* and *Ascl1* expression defines four NPC pools, that are uniformly distributed and ‘imbalanced’ in lissencephalic cortices (i.e. *Neurog2-*rich in rodents, *ASCL1*-rich in primate allocortex), and modular and ‘balanced’ in folded, and pseudo-folded cortices (Figure 7R).

## DISCUSSION

We set out to determine how a regular pattern of neurogenesis is sustained in the lissencephalic rodent brain by studying the proneural TFs Neurog2 and Ascl1, critical architects of the decision by NPCs to proliferate or differentiate (Wilkinson et al., 2013). Through the use of novel *in vivo* lineage tracing tools, we stratified cortical NPCs into four distinct pools that we showed have biased potentials towards neuronal (Neurog2^+^, double^+^), oligodendrocyte (Ascl1^+^, double^+^) and astrocyte (pro^−^) lineages. By combining *in vivo* lineage tracing with time lapse microscopy, we further found that double^+^ NPCs mark a lineage bifurcation point between neuronal and glial cell fates that is maintained through cross-repressive TF interactions. Compared to proneural single^+^ NPCs, double^+^ NPCs display several unique phenotypic properties, including: (1) more open chromatin and a more complex GRN that is conducive to bipotency, containing central Neurog2 and Ascl1 nodes and displaying dependency on both TFs; (2) a G_2_-pausing phenotype linked to regenerative competence in other species (Sutcu and Ricchetti, 2018), and (3) a unique role as Notch ligand-producing ‘niche’ cells that control neurogenic regularity to prevent cortical folding. Solidifying the importance of proneural genes in cortical folding, using RNA-Scope we showed that *NEUROG2* and *ASCL1* are expressed in a modular pattern in macaque cortex and pseudo-folded human brain organoids. While Notch signaling has previously been proposed to be important for brain folding based on modular expression patterns in ferret neocortex (de Juan Romero et al., 2015), and the expansion of a novel *NOTCH2NL* gene in human cortices (Fiddes et al., 2018), functional studies had not been performed. Our identification of proneural double^+^ NPCs as Notch ligand-expressing niche cells that are essential to maintain neurogenic continuity and to prevent cortical folding in the rodent cortex is a novel finding that provides the first functional support for the idea that discontinuous Notch signaling is associated with gyrencephaly.

How progenitor cells gradually become lineage restricted has been the subject of debate in many lineages, but nowhere has this question been more carefully dissected than in hematopoiesis (Brand and Morrissey, 2020). Here, several TF pairs have been identified that maintain bipotency through mutual cross-repression. A generalized computational model of stem and progenitor cell maintenance has been proposed that involves the balance of opposing differentiation forces, mediated by cross-repressive TF pairs, or lineage specifiers (Okawa et al., 2016). This model has gained experimental support from studies of embryonic stem cells (ESCs) (Dillon, 2012) and hematopoietic progenitors (Chickarmane et al., 2009), leading to a generalized concept in which antagonist TFs are coexpressed in bipotential progenitors, with changes in their relative levels pushing progenitors towards one fate or another. Notably, a recent study using Cytof has provided indisputable proof for this specification model in hematopoiesis (Palii et al., 2019). Here, using short-term and long-term split-cre lineage tracing systems, we similarly found that Neurog2 and Ascl1 are co-expressed, cross-repressive TFs that control a binary choice between two cell fates; neuronal and glial (oligodendrocyte), in cortical NPCs, providing the first evidence that a pair of antagonistic TFs also regulates cell fate decisions in the nervous system. The finding that *NEUROG2* and *ASCL1* expression levels are more ‘balanced’ in folded cortical structures suggests that tighter regulatory controls, achieved through TF co-expression and cross-repression, may be at play to control the timing and spacing of neurogenesis.

Cell fate decisions are not only dictated by the repertoire of TFs expressed, but also by the chromatin landscape. In neural lineages, Neurog2 and Ascl1 are pioneer factors that bind closed chromatin and facilitate opening of these sites (Aydin et al., 2019; Wapinski et al., 2013). Indeed, using an *in vitro* overexpression system in mouse embryonic stem cells, Ascl1 and Neurog2 were shown to bind to largely non-overlapping sites in the genome, and through their binding, establish distinct chromatin landscapes (Aydin et al., 2019). Our study provides in vivo support for the idea that Neurog2 and Ascl1 may participate in reshaping the chromatin landscape for subsequent fate determination, as the four NPC pools defined by *Neurog2* and *Ascl1* expression, we identified a continuum of lineage restriction based on the amount of open chromatin (i.e., hotspots), ordering double^+^, Ascl1^+^, pro^−^, and Neurog2^+^ NPCs from lowest to highest that correlates with their differentiation biases *in vitro* and *in vivo.* Notably, the lack of a complete restriction to one lineage or the other was expected as lineage biases are thought to gradually accumulate before ultimately culminating in cell fate choice and differentiation (Brand and Morrissey, 2020). Lineage biases of Neurog2^+^, Ascl1^+^ and pro^−^ NPCs towards neuronal, oligodendrocyte and astrocyte differentiation, respectively, was consistent with prior studies of individual genes (Li et al., 2014; Li et al., 2012; Parras et al., 2007), but the bipotency of double^+^ NPCs was an unexpected and new finding.

All pro^+^ NPCs had longer cell cycles than pro^−^ NPCs, which is generally thought to be conducive to differentiation, while conversely, increased pluripotency is associated with G_1_ and G_2_ shortening (Dalton, 2015). Notably, we also observed S phase lengthening in pro^+^ NPCs, similar to increased S-phase in Ascl1^+^ NPCs in the adult SVZ (Ponti et al., 2013). Global chromatin remodeling occurs in S-phase, and it is thus a period when cell fate restriction can occur (Ma et al., 2015). Indeed, in the cortex, heterochronic transplantations of NPCs in S-phase can change their laminar identities, whereas cell fates are no longer plastic after G_2_/M-phase (McConnell, 1995). In addition, we found evidence for G_2_-pausing in double^+^ NPCs, which is linked to stem cell maintenance in the zebrafish myotome, Hydra embryo and in ‘super-healer’ mice (Sutcu and Ricchetti, 2018), and may further contribute to the lack of lineage commitment of double^+^ NPCs. The idea is that cells paused in G_2_ have passed the energy-consuming effort of DNA replication and are thus poised in a flexible state where they can quickly undergo mitosis and divide into two daughter cells of different identities. Finally, double^+^ NPCs showed highest expression of *Cdkn1a,* a known G_2_ pausing factor (Bunz et al., 1998). Notably, *Cdkn1a* is essential for a long-term maintenance of NSCs that are associated with neural cell regeneration (Kippin et al., 2005), while *Cdkn1c* and *Trp73,* which are highly expressed in double^+^ NPCs, are also required for NSC maintenance (Furutachi et al., 2015; Talos et al., 2010).

Cortical folding has evolved independently several times during evolution (Lewitus et al., 2013). The major contributing force is the expansion of oRG, which generate more upper layer neurons to increase overall cortical mass, while also complexifying neuronal migratory routes to generate neuronal sparse and dense regions in the cortical plate (Del Toro et al., 2017). Several groups have induced cortical folding in mice, including by the knockdown of *Trnp1*, a nuclear protein (Stahl et al., 2013), overexpression of human-specific *ARHGAP11B* (Florio et al., 2015) or *TBC1D3* (Ju et al., 2016), or double knock-out of *Lmx1a;Lmx1b* (Chizhikov et al., 2019). Similarly, double knock-out of *Flrt1/Flrt3* adhesion molecules increases migration and tangential dispersion, leading to neuronal clustering and cortical folding in mice (Del Toro et al., 2017). These studies suggest that cortical folding is induced by oRG expansion, however, increasing the basal progenitor pool alone does not induce cortical folding in rodents (Nonaka-Kinoshita et al., 2013). Instead, folding requires a modular pattern of neurogenesis. Our study has revealed that the regular positioning of double^+^ NPCs expressing high levels of Notch ligands, which are interspersed with NPCs that express low levels, controls neurogenic patterning. We conclude that double^+^ NPCs act like ‘niche’ cells given their critical importance in maintaining neurogenic patterning. Interestingly, in the adult brain, Notch ligands (Jag1) expressed on endothelial cells instead act as niche cells to maintain NSC quiescence and prevent differentiation and depletion (Ottone et al., 2014). Future studies could address whether adult NPCs also co-express Neurog2 and Ascl1, and whether their role as niche cells is maintained until adulthood.

In summary, by characterizing Neurog2/Ascl1 single and double^+^ cortical NPCs, we have gained unprecedented new insights into how fate determination is controlled, and in so doing, we uncovered a novel role for double^+^ NPCs as essential for regular neurogenesis to promote a lissencephalic structure. This study has profound implications for our understanding not only of the regulatory measures that underlie the development of lissencephaly versus gyrencephaly, but also of the function of distinct NPC pools, and of proneural gene cooperativity.

## Acknowledgements

This work was supported by operating grants to CS from the Canadian Institutes of Health Research (MOP-44094, MOP-125905, PJT – 406690). CS holds the Dixon Family Chair in Ophthalmology Research. We thank Magdalena Gotz and Marjorie Brand for critical comments. SH was supported by Scholarships from the Cumming School of Medicine, Ontario Graduate Scholarship, UofT Vision Science Research Program, Peterborough K.M. HUNTER Charitable Foundation and Margaret and Howard GAMBLE Research Grant. GW was supported by Frederick Banting and Charles Best Canada Graduate Scholarship, Alberta Innovates Health Solutions scholarship, and ACHRI/CIHR Training Grant Studentship. SO was supported by an FNR CORE grant (C15/BM/10397420). RD was supported by a CIHR Canada HOPE Fellowship. LA was supported by an ACHRI/CIHR Training Grant Studentship.

## Author Contributions

SH: conceptualization, data curation, formal analysis, investigation, methodology, visualization, validation, writing – original draft, writing – review and editing

GAW: conceptualization, data curation, formal analysis, investigation, methodology, validation, writing – review and editing

SO: data curation, formal analysis, software, validation, visualization, writing-review and editing

LA: data curation, formal analysis, investigation

RD: data curation, formal analysis, investigation

IF: investigation, formal analysis, methodology, software

MB: formal analyses, software, validation

VCo: formal analysis, investigation

VCh: formal analysis, investigation

DZ: formal analysis, investigation

SL: formal analysis, investigation

JG: investigation

FM: investigation

YT: investigation

VEA: formal analyses, software

AMO: formal analysis, investigation

ER: investigation

YI: investigation

JWK: investigation

WW: investigation

WR: investigation

IK: resources, supervision

JAC: resources, supervision

DK: resources, supervision

DSC: resources, formal analyses, software

CD: resources, supervision

AS: resources, supervision

JB: resources, supervision, validation, writing – review and editing

AdS: funding acquisition, resources, supervision, validation, writing – review and editing

CS: funding acquisition, conceptualization, project administration, resources, supervision, validation, writing – original draft; writing – review and editing

## STAR Methods

### Key Resources Table

### Contact for Reagent and Resource Sharing

Further information and request for resources and reagents should be directed to and will be fulfilled by the Lead Contact, Carol Schuurmans.

### Experimental Models and Subject Details

None of the animals used in our experiments had been previously used for other procedures. The animals presented a healthy status and were employed independently of their gender. The developmental stage of experimental models was chosen depending on the requirements of each experiment, as further detailed below.

#### MICE

##### Animal sources and maintenance

Animal procedures were approved by the University of Calgary Animal Care Committee (AC11-0053) and later by the Sunnybrook Research Institute (16-606) in compliance with the Guidelines of the Canadian Council of Animal Care. The generation of *Ascl1^GFPKI^* (Leung et al., 2007) (Jackson Lab: 012881) and *Neurog2^GFP^*^KI^ (Britz et al., 2006) null mutant mice were previously described. Rosa-DTA (Jackson Lab: 009669), Rosa-tdtomato, and Rosa-zsGreen (Jackson Lab : 007906) mice were purchased from Jackson Laboratory. The generation of new transgenic mouse lines (*Neurog2^Flag-mCherry^*^KI^, *Neurog2^N-Cre^*^KI,^ *Ascl1^C-Cre^*^KI^) are described below. All animals were maintained on a CD1 background, and wild-type CD1 mice were used in all non-transgenic experiments. Analyses were performed between E10.5-E17.5 and at P0-P7, as outlined in the text. Sex was not considered due to the difficulty in assigning sex at these stages.

##### Generation of transgenic mice

<u>*Neurog2^Flag-mCherry^*^KI/+^ mice</u> were generated by homologous recombination in embryonic stem cells (ESCs). A targeting vector (pPNT-Neurog2Flag-IRESmCherry-LPN) was linearized by *Swa*I and electroporated in G4 (129xC57BL/6) ES cells. Transfected ES cells were exposed to positive-negative selection with G418/gancyclovir. Genomic DNA samples were digested with *Not*I/*Spe*I and probed with 5’ (1.5-kb *Not*I/*Kpn*I) and 3’ (1-kb *Eco*RI/*Spe*I) external probes (Fode et al., 1998). For wild-type DNA, the 5’ and 3’ probes recognized the same 18.3-kb fragment whereas, for mutant DNA, an 8.3-kb 5’ and 11.6-kb 3’ fragment were recognized. The U of Calgary Transgenic Services aggregated cell lines identified as positive with morulae to produce chimeras, which were then provided to us for further analysis. Chimeras were bred with CD1 mice and agouti offspring genotyped by PCR using primers *Neurog2Flag-mCherry*\*F and *Neurog2Flag-mCherry*\*R, which gave rise to a 413-bp amplicon when the mutant allele was present. 35 cycles of 98°C/1 sec and 60°C/30 sec using primers for wild-type (*Neurog2*\*F and *Neurog2*\*R) and 95°C/5 min plus 40 cycles of 95°C/1 min, 60°C/1 min and 72 °C/1 min for mutant (*Neurog2Flag-mCherry*\*F and *Neurog2Flag-mCherry*\*R) alleles. *Neurog2*\*F: 5’ TAGACGCAGTGAC TTCTGTGACCG 3’. *Neurog2*\*R: 5’ ACCTCCTCTT CCTCCTTCAACTCC 3’. *Neurog2Flag-mCherry*\*F: 5’ ACAAACAACGTCTGTAGCGACCCT. 3’ *Neurog2Flag-mCherry*\*R: 5’ CACCTTGAAGCGCATGAACTCCTT 3’.

*Neurog2^N-Cre^*^KI^ and *Ascl1^C-Cre^*^KI^ transgenics were generated by Cyagen Biosciences (https://www.cyagen.com/us/en/) using homologous recombination in ESCs. Targeting strategies replaced genomic sequence between the *Neurog2* or *Ascl1* START and STOP codon with Ncre (amino acids 19-59) or Ccre (amino acids 60-343) modified with a nuclear-localization signal, GCN4 coiled-coil domain, flexible linker, and cre-recombinase sequences (for *Neurog2* and 60-343 for *Ascl1*) (Beckervordersandforth et al., 2010; Hirrlinger et al., 2009). Targeting vectors were cloned by VectorBuilder (https://en.vectorbuilder.com) with (109-FUV-hGFAP-NCre or 106-FUV-P2-CCre, respectively).

#### NON-HUMAN PRIMATES

##### Animal sources and maintenance

The use of cynomolgus monkey (Macaca fascicularis) in this study and all experimental protocols were approved by the Animal Care and Use Committee (CELYNE; protocols C2EA42-12-11-0402-003 and APAFIS#3183). Surgical procedures and animal experimentation were in accordance with European requirements 2010/63/UE. Animals were housed in a controlled environment (temperature: 22 ± 1°C) with 12 hr light/12 hr dark cycle (lights on at 08:00 a.m.). All animals were given commercial monkey diet twice a day with tap water ad libitum and were fed fruits and vegetables once daily. During and after experiments, monkeys have been under careful veterinary oversight to ensure good health. Foetuses from timed-pregnant cynomolgus monkeys (Embryonic day 73-E73) were delivered by caesarean section as described (Lukaszewicz et al., 2005).

#### HUMAN CEREBRAL ORGANOIDS

##### Generation of human cerebral organoids (COs)

Feeder-free H1 hESCs (WiCell) were cultured on Matrigel in TeSR™-E8™ kit for hESC/hiPSC maintenance (StemCell Tech; #05990). hESCs were used to generate COs using the STEMdiff Cerebral Organoid Kit (StemCell Tech; #08570) and STEMdiff Cerebral Organoid Maturation Kit (StemCell Tech; #08571) as directed by the manufacturer. Culture of human ESCs received approval from the Canadian Institutes of Health Research (CIHR) Stem Cell Oversight Committee (SCOC) to CS and was approved by the Sunnybrook REB (PIN: 264-2018).

### Method Details

#### Immunohistochemistry

Brains were dissected in phosphate-buffered saline (PBS) and fixed in 4% paraformaldehyde (PFA)/1X PBS overnight at 4°C. Brains were washed three times in 1X PBS, cryoprotected in 20% sucrose/1X PBS overnight at 4°C, blocked in OCT, and cryosectioned at 10 μm. Immunohistochemistry was performed as described (Li et al., 2014). Primary antibodies used included goat anti-Neurog2 (1:500, Santa Cruz Biotechnology #sc-19233), rabbit anti-Neurog2 (1:500, Invitrogen #PA5-78556), mouse anti-Ascl1 (1:100, BD Biosciences #556604), rabbit anti-GFP (1:500, Invitrogen #A-11122), goat anti-GFP (1:500, Abcam #ab5449), goat anti-mCherry (also detects tdTomato; 1:500, Sicgen #AB0040), rat anti-mCherry (1:500, Invitrogen #M11217), rabbit anti-NeuN (1:500, Abcam #ab177487), mouse anti-NeuN (1:100, Chemicon #MAB377), rabbit anti-Ki67 (1:500, Abcam #16667), rat anti-BrdU (1:50, BioRad #OBT0030S), mouse anti-BrdU (Mobu-1; 1:50, Invitrogen #B35128), mouse anti-Tuj1 (β3-tubulin, 1:500, BioLegend #801202), goat anti-Pdgfrα (1/500, R&D Systems #AF1062), rat anti-Pdgfrα (1:500, eBioscence #14-1401-82), rabbit anti-GFAP (1:500, DakoCytomation #Z0334), rat anti-GFAP antibody (1/500, Thermo Fisher Scientific #13-0300), rabbit anti-Sox9 (1:500, Millipore #AB5535), rabbit anti-Pax6 (1:350, Cedarlane #PRB-278P), rabbit anti-Tbr2 (1:500, Abcam #ab23345), rabbit anti-zsGreen (1:500, Takara #632474), rabbit anti-Tbr1 (1:500, Abcam #Ab31940), mouse anti-Satb2 (1:500, Abcam #ab51502), mouse anti-Nestin (1:250, Millipore #MAB353), chicken anti-GFP (1:500, Abcam#ab13970) and rabbit anti-S100b (1:100, Dako/Agilent #Z031129-2). Secondary antibodies were all diluted to 1/500 and conjugated to different fluorophores, including Cy3 (red: mouse IgG, Jackson Immunoresearch #715-166-150), Alexa Fluor™ 405 (blue: mouse IgG, Abcam #Ab175659), Alexa Fluor™ 488 (green: rabbit IgG #A21206; goat IgG #A11055; rat IgG #A21208; mouse IgG #A11029, mouse IgG1 #A21121; chicken IgG #A11039, all from Invitrogen), Alexa Fluor™ 568 (red: rabbit IgG #A10042; goat IgG #A11057; rat IgG #A11077; mouse IgG #A11004; mouse IgG1 #A21124) or Alexa Fluor™ 647 (far-red: rabbit IgG, Invitrogen #A31573; rat IgG, Jackson ImmunoResearch #712-605-153).

#### RNAscope assay

The RNAscope^®^ Multiplex Fluorescent Detection Kit v2 (ACD #323110) was used according to the manufacturer’s directions. Briefly, cryosections were post-fixed with 4% PFA for 15 min at 4°C and then dehydrated with serial incubations in 50%, 70% and 100% EtOH for 5 min each at room temperature. Sections were then incubated in hydrogen peroxide solution for 10 min at room temperature, in 1x target retrieval solution for 5 min at 95°C, and washed with distilled water. Protease Plus was then added for 15 min at 40°C and removed with washing buffer. RNA probes were then applied for 2 hrs at 40°C as indicated. Probes included Mm-*Ascl1* (#313291), Mm-*Neurog2-C2* (#417291-C2), Mm-*Hes1* (#417701), Mfa-*ASCL1* (#546591), Mfa-*NEUROG2*-C2 (#546589-C2), Hs-*ASCL1* (#459728) and Hs-*NEUROG2*-C2 (#546601-C2). Amplification and staining steps were performed according to manufacturer’s instructions. Opal^TM^ 570 (Akoya #FP1488001KT; 1:1500) was applied for channel 1 and Opal^TM^ 520 (Akoya #FP1487001KT; 1:1500) was applied for channel 2.

#### BrdU staining and cell cycle analyses

Bromodeoxyuridine (BrdU; Sigma, Oakville, ON) was injected IP at 100 µg/g body weight 30 min or 24 hr before dissection as indicated. To measure cell cycle length, BrdU were administered every three hours up to 15h in E12.5 pregnant dams. Mice cortices were dissected 30 min, 1, 2, 3, 6, 9 and 15 h after BrdU injection. For BrdU co-immunolabeling, sections were first immunolabeled with the other antibody (as indicated), post-fixed with 4% PFA for 10 min and washed 3 times by PBS with 0.1% Triton X-100. Sections were treated with 2N HCl for 25 min at 37°C for antigen retrieval, and immunolabeled with anti-BrdU.

#### Fluorescence activated cell sorting (FACS) and flow cytometry

Brains from E12.5 *Neurog2*^Flag-^ *^mcherry^*^KI/+^; *Ascl1^GFP^*^KI/+^ embryos were dissected in PBS and separated by FACs according to GFP and mCherry expression. Double KI cortices were digested in PBS (Ca^++^/Mg^++^ free) containing 25 μg/ml trypsin (Sigma #T1005-1G) for 10 minutes at 37°C. Digestion was stopped with 20% FBS and cells were triturated ∼40 times. Cells were resuspended in PBS containing 5 mM EDTA and 0.1% BSA. Dissociated cells (2.5 µl/1×10^6^ cells) were stained with Viability Dye eFluor 780 (eBioscience #65-0865-14) and with anti-CD15-Alexa Fluor 647 (BD Bioscience #560120) or anti-CD133-APC (eBioscience #17-1331-81) and PerCP-eFluor 710 (eBioscience #46-1331-82) according to the manufacturer’s directions. For cell cycle analysis, cells were also stained with Hoechst 33342 (Thermo Fisher Scientific #62249) according to the manufacturer’s protocol. FACS and cell cycle analysis was performed with a BD FACS Aria III cell sorting system. Quantitation of cell cycle phases was performed with FlowJo software using a Dean Jett Fox algorithm.

#### Neurosphere assay

FACS sorted cells were seeded in 0.2% gelatin coated 24-well plates at clonal density (5,000 cells/well) and cultured for 7 days in Neurosphere media containing DMEM/F12 (3:1), human FGF2 (40 ng/mL), human EGF (20 ng/mL), B27 supplement minus vitamin A (2%), Penicillin/streptomycin (0.1%), Fungizone (40 ng/mL), and cyclopamine (0.5 µg/ml). Primary neurospheres were counted and photographed using an AxioVision program (Carl Zeiss).

#### Directed differentiation assay

FACS sorted cells were plated in 8 well chamber slides coated with poly-L-ornithine and laminin for the differentiation of neurons and oligodendrocytes, or Geltrex® matrix (Thermo Fisher Scientific #12760) for the differentiation of astrocytes. Cells were incubated for 1 day in Stem cell media, containing KnockOut^tm^ D-MEM/F12, GlutaMax^tm^-I supplement (2 mM), bFGF (20 ng/ml), EGF (20 ng/ml), 2% StemPro® Neural Supplement, Penicillin/streptomycin (0.1%) and Fungizone (40 ng/mL). To induce neuronal differentiation, cells were grown in Neurobasal® medium, 2% B-27® Serum-Free Supplement (Thermo Fisher Scientific #17504) and GlutaMax^tm^-I supplement (2 mM). Astrocyte differentiation medium contained D-MEM, 1% N-2 Supplement (Thermo Fisher Scientific #17502), GlutaMax^tm^-I supplement (2 mM) and 1% FBS. Oligodendrocyte differentiation medium contained Neurobasal® medium, 2% B-27® Serum-Free Supplement (Thermo Fisher Scientific #17504), GlutaMax^tm^-I supplement (2 mM) and T3 (Sigma #D6397). Cells were fed every 2 days with 2X media for 4 DIV or 10 DIV. At experimental endpoint, cells were fixed with 4% PFA for 15 min at room temperature and immunostained using mouse anti-Tuj1 (neuronal III β-tubulin, 1/500, Covance, Laval, QC, #801202), rat anti-Pdgfrα (1/500, BD Pharmingen™, #558774) and rat anti-GFAP (1/500, Thermo Fisher Scientific #13-0300). Secondary antibodies were conjugated to Alexa Fluor™ 568 (as indicated).

#### Time lapse imaging

Cortices from E14.5 *Neurog2^Flag-mCherry^*^KI^;*Ascl1^GFP^*^KI/+^ embryos were cut to 300 μm in ice cold DMEM/F12 using a Leica VT1200S vibratome. Slices were immersed in ice cold type 1a collagen (Cellmatrix, Nitta Gelatin, cat. 631-00651) on 30 mm glass cover slips which were then incubated at 37°C in a Zeiss chamber system filled with 40% O_2_, 5% CO_2_, and 55% N_2_. After a 30 min incubation, slices were cultured in 2-3 mL of slice culture medium as previously described (Shitamukai et al., 2011). Images for the dorsal telencephalon were obtained on an inverted confocal microscope (Zeiss) with either a 20X or 40X objective. Pictures were taken every 10 minutes in ∼20 μm Z stack sections with 8-10 images per stack. Using an automated platform, multiple regions on the same slice were imaged over an ∼18-hour period. Time lapse images were further processed and analyzed with Fiji software.

#### Luciferase Assays

P19 embryonic carcinoma cells (ATCC# CRL-1825) were maintained in Minimum Essential α Medium (Gibco) supplemented with 10% fetal bovine serum and 50 units/ml penicillin-streptomycin (Gibco). P19 cells were seeded into 24-well plates (Nalge Nunc) 24 hr before transfection. The p*NeuroD*^1kb^ (Huang et al., 2000), *Rnd2* (Heng et al., 2008)*, Sox9* (Li et al., 2014) and *Dll1* (Castro et al., 2006) *luciferase* reporters were previously described. Transfections were performed using Lipofectamine Plus reagent (Invitrogen), following the manufacturer’s protocol, with 0.1 μg of each plasmid and 0.15 μg of a Renilla plasmid (transfection control). 4-6 hr post-transfection, Opti-MEM media (Gibco) was replaced with fresh media. 24 hr later, cells were harvested and firefly luciferase and Renilla luciferase activities were measured using the Dual-luciferase Reporter Assay System (Promega #E1910) following the manufacturer’s instructions. Measurements were made using a TD 20/20 Luminometer (Turner Designs). For the analysis of *luciferase*/*Renilla* assays, *luciferase* data was normalized by dividing raw light readings by the corresponding Renilla values.

#### Immunoprecipitation

Immunoprecipitation (IP) was performed as described (Han et al., 2018). Briefly, micro-dissected E12.5 *Neurog2^FLAG-mCherryKI/+^* cortices or NIH-3T3 (ATCC CRL-1658) cells transfected with pCIG2-*Ascl1* and either pCS108-FLAG or pCS108-*Neurog2*-FLAG (Han et al., 2018) expression vectors were harvested after 48 hr and lysed in NET2 lysis buffer (0.05% NP40, 150 mM NaCl, 50 mM Tris-Cl, pH 7.4) containing protease (1X protease inhibitor complete, 1 mM PMSF), proteasome (7.5 μM MG132) and phosphatase (50 mM NaF, 1 mM NaOV_3_) inhibitors. 400 μg of protein lysate was immunoprecipitated using anti-FLAG M2 beads (Sigma) overnight at 4°C. The sample was divided in two and one half was incubated with DNaseI (2 U/ml; Ambion). FLAG-beads were washed 5 times in lysis buffer, resuspended in SDS-PAGE loading dye, and run on 8% or 10% SDS-PAGE gels followed by Western blot analysis.

#### Western blot

Western blots were performed as described (Li et al., 2012) with: mouse-anti-Ascl1 (1:10,000, BD Biosciences #556604), rabbit anti-FLAG (1:10,000, Cell signaling #2368), rabbit anti-GAPDH (1:10,000, Cell signaling #2118), rabbit anti-Notch (Cleaved) (NICD, 1:1000, Cell signaling #4147). Western blot signals were converted to a chemiluminescent signal using an ECL kit (EG Healthcare) following the manufacturer’s instructions and visualized using X-ray film or a Bio-rad gel doc with GelCapture MicroChemi 2.2.0.0 software.

#### In utero electroporation

cDNA expression vectors were introduced into dorsal telencephalic (cortical) progenitors using i*n utero* electroporation as previously described (Dixit et al., 2011; Li et al., 2012). pCIG2-*Neurog2* (Mattar et al., 2008), pCIG2-*Ascl1* and pCIG2-*Neurog2*∼*Neurog2* (Li et al., 2012) expression vectors were described previously. To generate a *Neurog2∼Ascl1* tethered construct, the following oligos were annealed to make a tether, leaving *Pst*I ends for ligation: 5’-GGG GGT TCC GGC GGG GGT TCT GGA GGT GGG AGC GGC GGA GGG TCC GGC GGA GGA ACT GCA-3’; 5’GTT CCT CCG CCG GAC CCT CCG CCG CTC CCA CCT CCA GAA CCC CCG CCG GAA CCC CCT GCA-3’.

#### Proximity Ligation Assay (PLA)

P19 embryonic carcinoma cells were transfected with plasmids as indicated using Lipofectamine 2000 according to the manufacturer’s instructions. After 48 hrs, cells were fixed with 4% PFA for 15 min at room temperature and washed twice with PBS. The PLA assay was performed using a Duolink™ In Situ Red Starter Kit Mouse/Goat (Sigma Aldrich #DUO92103) following the manufacturer’s instructions. Primary antibodies included: anti-Neurog2 (1:200, Santa Cruz Biotechnology #sc-19233), mouse anti-Ascl1 (1:100, BD Biosciences #556604).

#### RNA-seq

RNA was extracted from cells using MagMAX™-96 Total RNA Isolation Kit (Invitrogen, #AM1830) according to the manufacturer’s protocol. Strand-specific mRNA sequencing libraries were constructed from 100 ng of total RNA using the TruSeq Stranded mRNA Library Prep Kit (Illumina, San Diego, CA, #20020594) and 101 base paired-end sequence reads were generated on the HiSeq 2500 platform (Illumina). Base calls were generated with RTA v1.18.64 software and pass filter reads were kept for further analysis. Fastq adapter trimming, quality assessment, alignment, and transcript quantitation were performed using assembly GRCm38.p6 and Ensembl v94 annotation as previously described (Kaewkhaw et al., 2015). Transcripts were retained for further analysis if they expressed >1.0 fragments per kilobase of exon model per million reads (FPKM from eXpress output) in all replicates of any group. Effective counts from the eXpress output of transcripts passing FPKM filtering were TMM normalized with edgeR v3.22.5 (Robinson et al., 2010). Quantitation of aligned reads to transcriptome and gene level summarization were achieved using Kallisto v0.44.0 for transcript isoform read assignment, isoform de-convolution, and quantitation and tximport v1.8.0 for gene level quantitation. We used the high-performance computational capabilities of the Biowulf Linux cluster at NIH (http://biowulf.nih.gov). Heatmaps with Euclidian clustering were generated using Heatmapper (Babicki et al., 2016). GO enrichment analysis was performed by EnrichR (Kuleshov et al., 2016).

#### Nanostring analysis

Total RNA from FACS sorted cells was isolated with Trizol reagent using the manufacturer’s instructions (Life Technologies). RNA integrity and concentration were measured with an Agilent 2100 bioanalyzer. RNA was hybridized to a custom made Nanostring CodeSet (Supplemental Table 1) and barcodes were counted on an nCounter digital analyzer using the manufacturer’s instructions (NanoString Technologies Inc.). Gene expression analyses were performed with nSolver Analysis Software. Gene expression was normalized relative to six spiked positive controls; three reference genes *Actb, Gapdh, Tubb*; and six negative controls to subtract background hybridization.

#### RT-qPCR

RNA was extracted from cells using Trizol reagent (Thermo Fisher Scientific #15596-026) or RNeasy Micro Kit (Qiagen, #74004) according to the manufacturer’s protocol. cDNA was synthesized with a RT^2^ First strand kit (Qiagen, cat. 330401) and qPCR was performed with RT^2^ SYBR Green (Qiagen, cat. 330500), both using the manufacturer’s protocols. Pre-validated RT^2^ qPCR primers were from Qiagen as follows: *Neurog2* (PPM28944A), *Ascl1* (PPM31367F), *Cdkn1a* (PPM02901B), *Cdkn1c* (PPM02895B), *Trp73* (PPM03436B), *Nestin* (PPM04735A), *Pax6* (PPM04498B), *Tbr2* (PPM32970F), *NeuN* (PPM60749A), *Gapdh* (PPM02946E), *B2m* (PPM03562A) and *Hprt* (PPM03559F). qPCR was performed with three biological replicates (N; RNA from three embryos) and three technical replicates per sample (n). Relative gene expression was determined using the delta-delta Cq method standardizing relative to reference genes (*Gapdh, B2m, Hprt)* and normalizing to pro^−^ NPC values (set at 1).

#### Nucleofection

Nucleofection was performed with a Lonza mouse neural stem cell kit according to the manufacturer’s instructions (Lonza #VPG-1004). Nucleofections were performed with ∼5 million cells dissociated from pooled E12.5 dorsal telencephalons. Nucleofected cells were then seeded for a Neurosphere assay as described except nucleofected cells were cultured for 10 days. To overexpress *Cdkn1a, Cdkn1c* and *Trp73,* cDNAs were cloned into the PiggyBAC Transposon vector (System Biosciences, Inc. #PB530A-2) containing a modified CAG promoter.

#### ATAC-seq

CD15^+^ GFP/mCherry negative, single, double positive cortical NPCs were dissociated from E12.5 *Neurog2^Flag-mCherry^*^KI/+^;*Ascl1^GFP^*^KI/+^ dorsal telencephalic tissue. ATAC-seq was performed as previously described, using 50,000 cells as input material for the transposase reaction with Nextera DNA library preparation Kit (Cat. No. FC-121-1030) (Buenrostro et al., 2015). Sequencing was performed with three biological replicates to a depth of approximately ∼330 million reads to delineate TF footprints. Heatmaps and average plots for the ATAC-seq reads near TSS were generated by NGSPLOT tool after merging the replicates of samples into a single BAM file and eliminating the duplications of reads. To identify bona fide open regulatory regions (i.e., true regions of local enrichment relative to the background ATAC-seq signal), we employed a commonly used hotspot (HS) algorithm with a FDR < 0.01 (John et al., 2011). We further refined our analysis by only examining HSs present in all three biological replicates, yielding one high confidence data set. Differentially accessible sites (DASs) were identified using statistical routines implemented in DiffBind Bioconductor package. Merging function was used to find all overlapping HSs and derive a single unique set of intervals for all samples to apply normalization and statistical testing of edgeR Bioconductor package. Each consensus interval in the Diffbind analysis was annotated with the distance to the nearest promoter and corresponding gene id was added using ChIPpeakAnno package. To predict which TF were binding to the unique regulatory regions in each cell type, we first used the Wellington algorithm to identify TF footprints within the ATAC-seq peaks (Piper et al., 2013), then used the Homer annotate peaks tool to identify known TF motifs (Heinz et al., 2010).

#### RNA in situ hybridization

RNA *in situ* hybridization was performed as described (Li et al., 2014; Li et al., 2012). The following templates that were used to generate anti-sense riboprobes: *Nfia* (*Kpn*I, T7), *Zfp423* (GenBank: BC079586, *Sal*I, T3), *Klf13* (GenBank: AK155298, *Sac*I, T3), *Foxp4* (GenBank: BC052407, *Sal*I, T3).

#### Gene regulation network analysis (GRN) and in silico perturbation assay

GRN analysis was performed using the program codes from (Okawa et al., 2015). Briefly, GRNs for pro^−^, Neurog2^+^, Ascl1^+^ and double^+^ NPCs were built using population specific TFs that were differentially expressed in each population with respect to the proneural negative population. GRN for proneural negative NPC was built using the union of differentially expressed TFs against the other three NPCs. Neurog2 and Ascl1 targets were identified using E14.5 telencephalon Neurog2 ChIP-seq data (Sessa et al., 2017) and Ascl1 ChIP-seq data provided by Dr. Diogo S. Castro. Transcriptional interactions were obtained as the union of the two ChIP-seq data and the MetaCore database. Interactions were removed if the chromatin regions of target genes were not open in all ATAC-seq replicates. After building GRN, *in silico* perturbation assays were performed by repressing the expression of either Neurog2 or Ascl1, or both together. The GRN simulation was carried out by the Boolean network formalism using the inhibitor dominant logic rule. When GRN reached a steady state, repressed genes were identified.

#### TUNEL assay

Cryosections were washed in PBS for 5 min, fixed with 4% PFA for 15 min at 37°C, and then permeabilized by Proteas K solution provided by a Click-iT® Plus TUNEL Assay kit (Invitrogen #C10619). Sections were washed in PBS for 5 min, fixed again in 4% PFA for 5 min at 37°C, rinsed with deionized water and subjected to Click-iT® Plus reaction with Alexa Fluor® 647 dye using Click-iT® Plus TUNEL Assay kit (Invitrogen #C10619) according to the manufacturer’s instructions. After Click-iT® Plus reaction, brain sections were stained with DAPI (Invitrogen #D1306) and mounted using Aqua-Poly/Mount (Polysciences #18606).

#### Imaging and image processing methods

Images of stained sections and cells were obtained using a Leica DMI8 fluorescent microscopy or a Zeiss Axiovert 200M confocal microscopy. Immunolabeled cells or RNAscope signals were counted using CellProfiler and Fiji software. Line plots for RNAscope signals were generated by the Plot profile function of Fiji software. Distance map analysis for *Hes1* RNAscope signals was performed using the Geometry to Distance Map function with 150 threshold and inverse options of Fiji software. The mean value from Geometry to Distance Map analysis was calculated and compared between the groups.

### Quantification and statistical analysis

All experiments used for statistical analysis were performed a minimum of three times (biological repeats from individual embryos and independent experiments, N-values are indicated in the figure legends). Individual biological replicates are shown as individual dots in the plots after averaging technical replicates. All statistical analysis was performed as described in the legend using GraphPad Prism Software version 8.0 (GraphPad Software). Unpaired two tailed Student’s t test was used to calculate statistical significance between two experimental groups. For multiple comparison between more than two experimental groups, one-way ANOVA with a Tukey post-hoc analysis was used. Error bars represent standard error of the mean (SEM). Significance was defined when p-value is less than 0.05.

## Data and software availability

RNA-seq data submitted to GEO: https://www.ncbi.nlm.nih.gov/geo/query/acc.cgi?acc=GSE151775

ATAC-seq data submitted to GEO: http://www.ncbi.nlm.nih.gov/geo/query/acc.cgi?acc=GSE84120

Uncropped western blot images deposited in Mendeley data: http://dx.doi.org/10.17632/wgpsd9yyk2.1

---

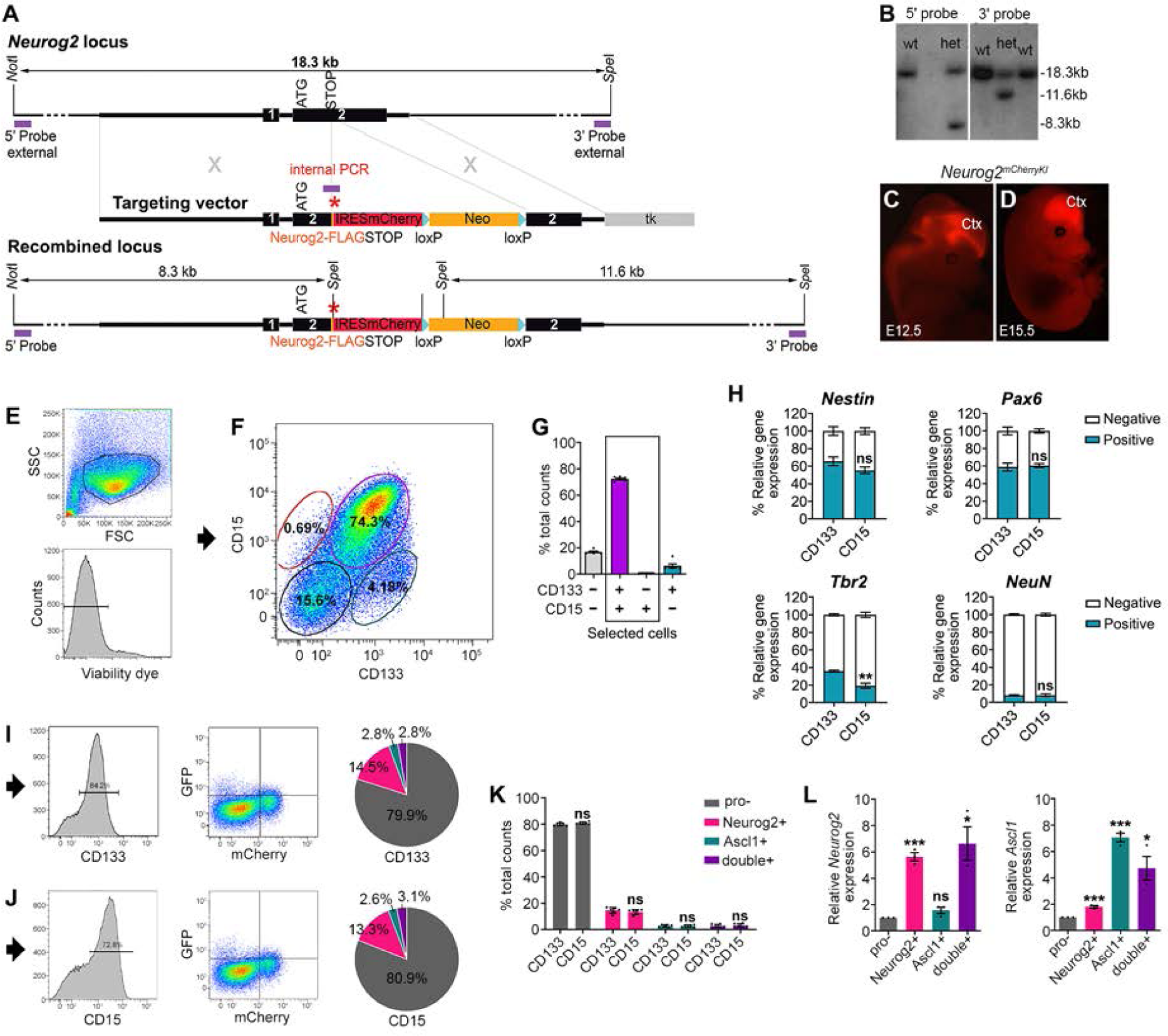

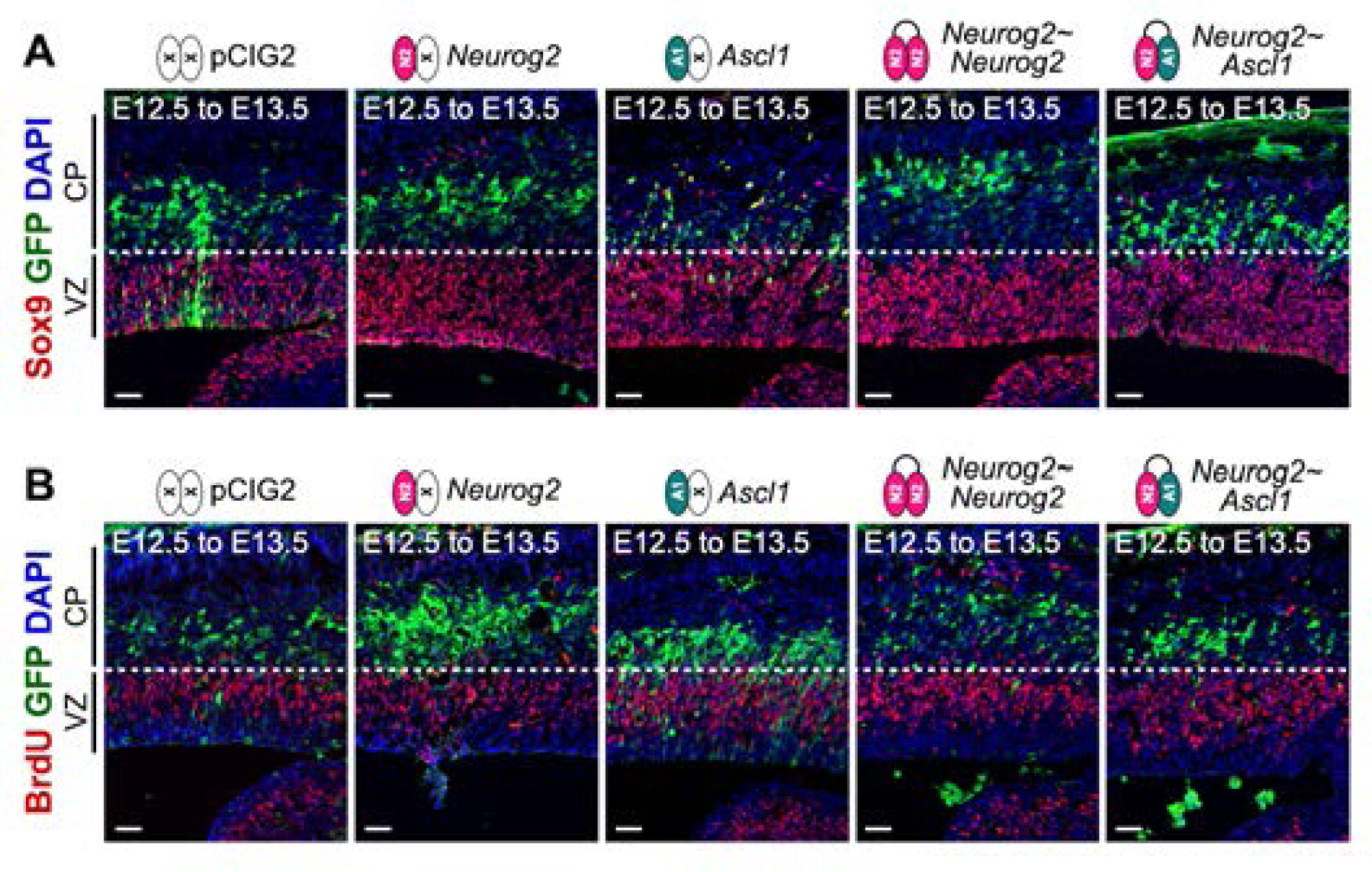

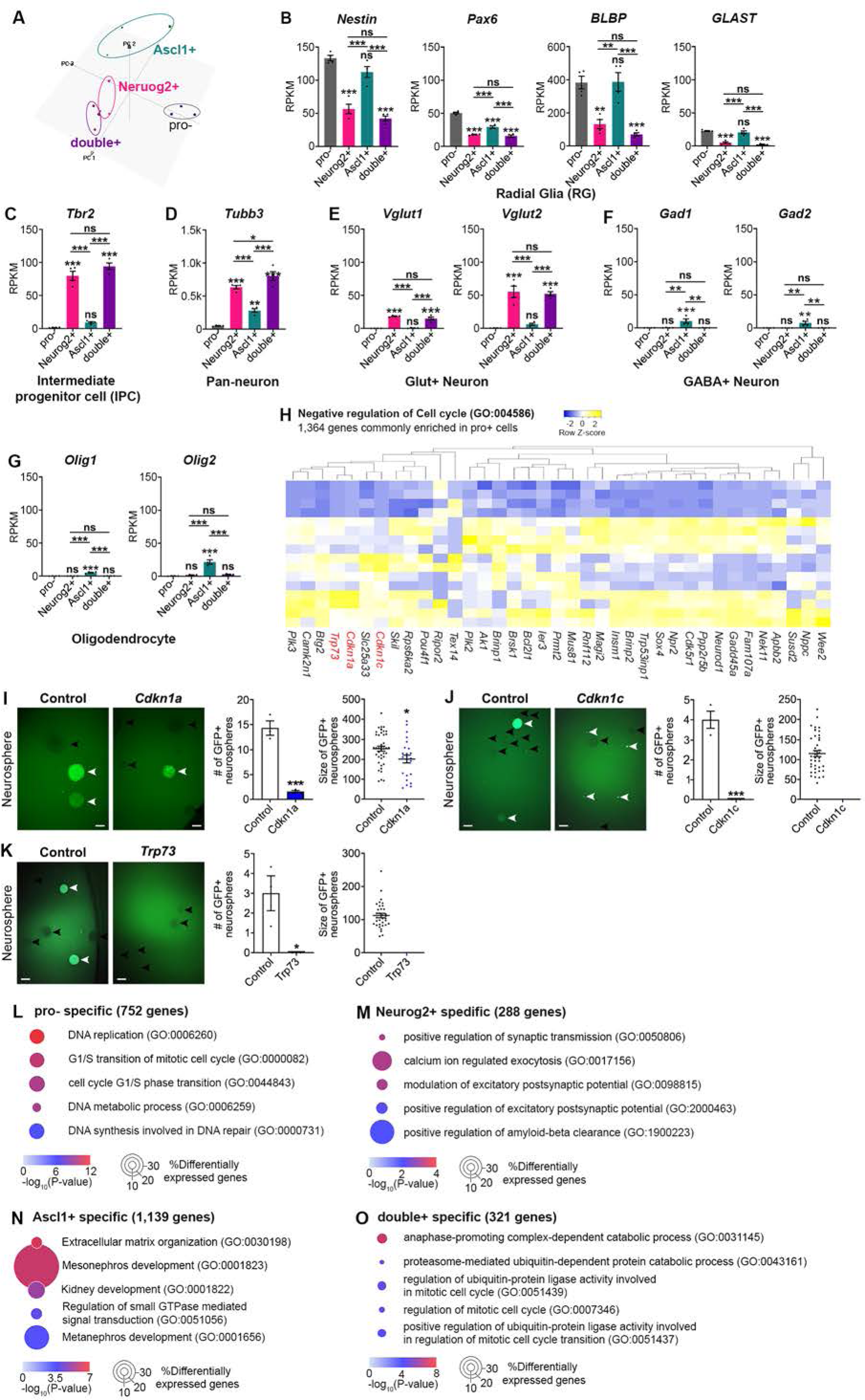

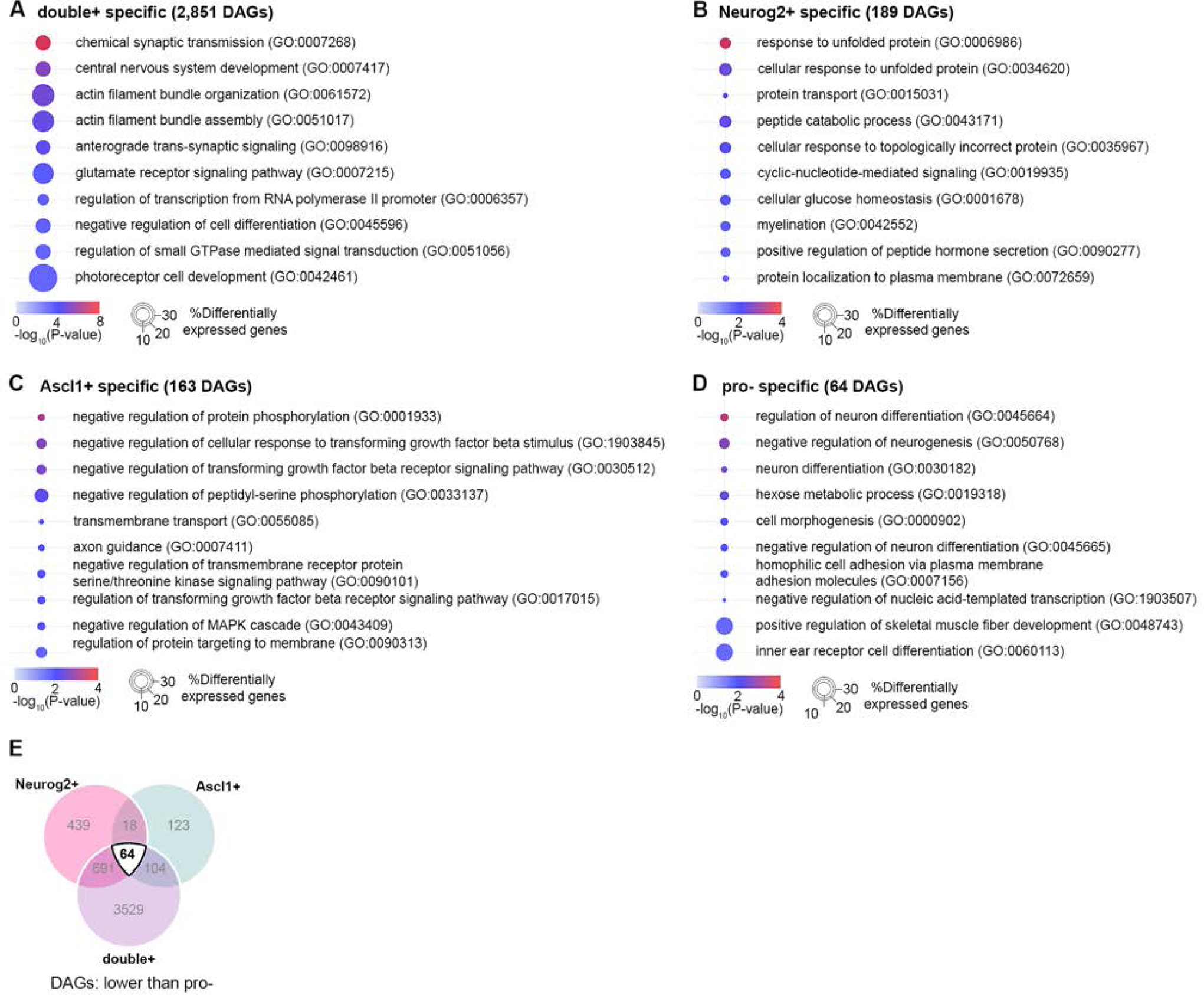

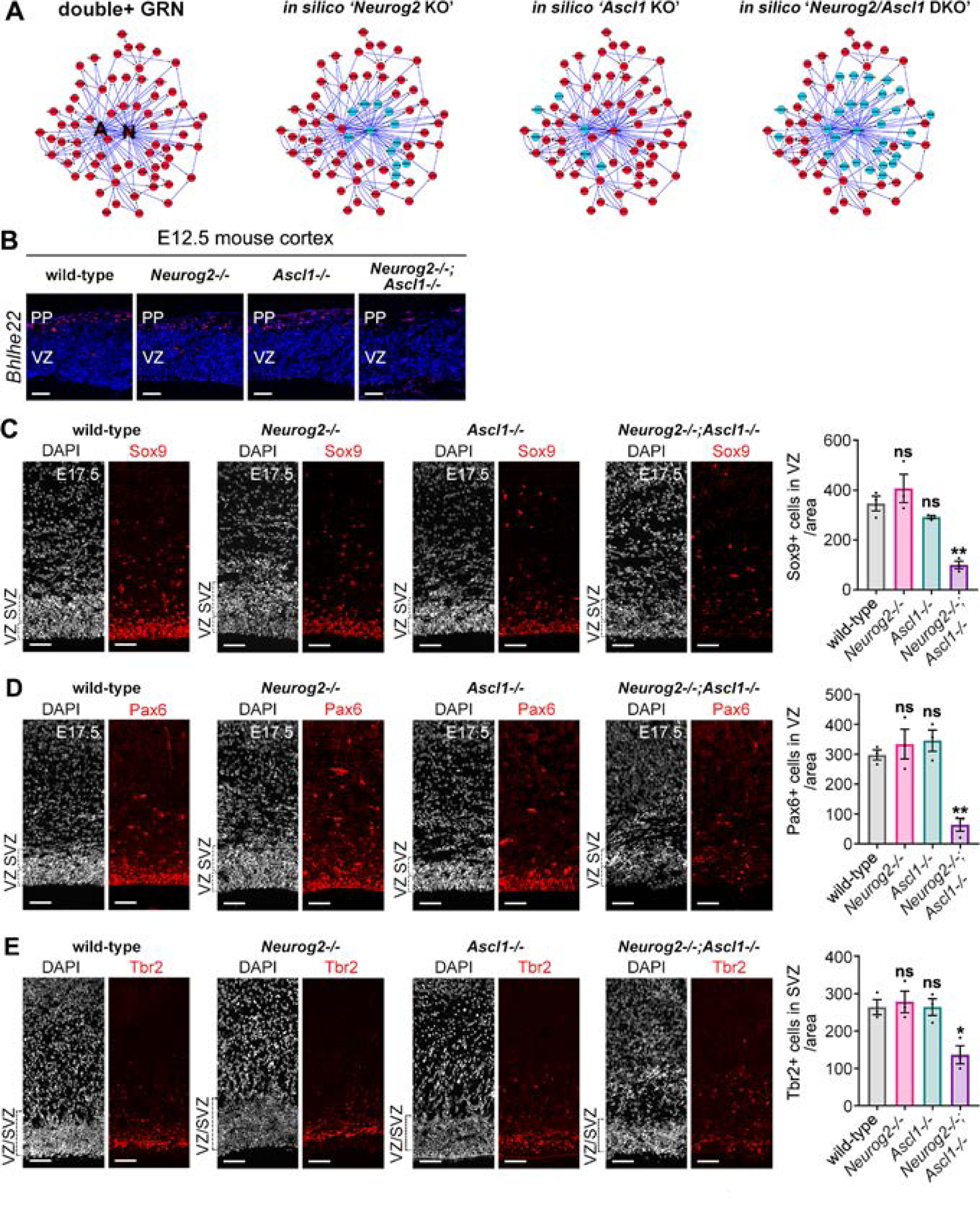

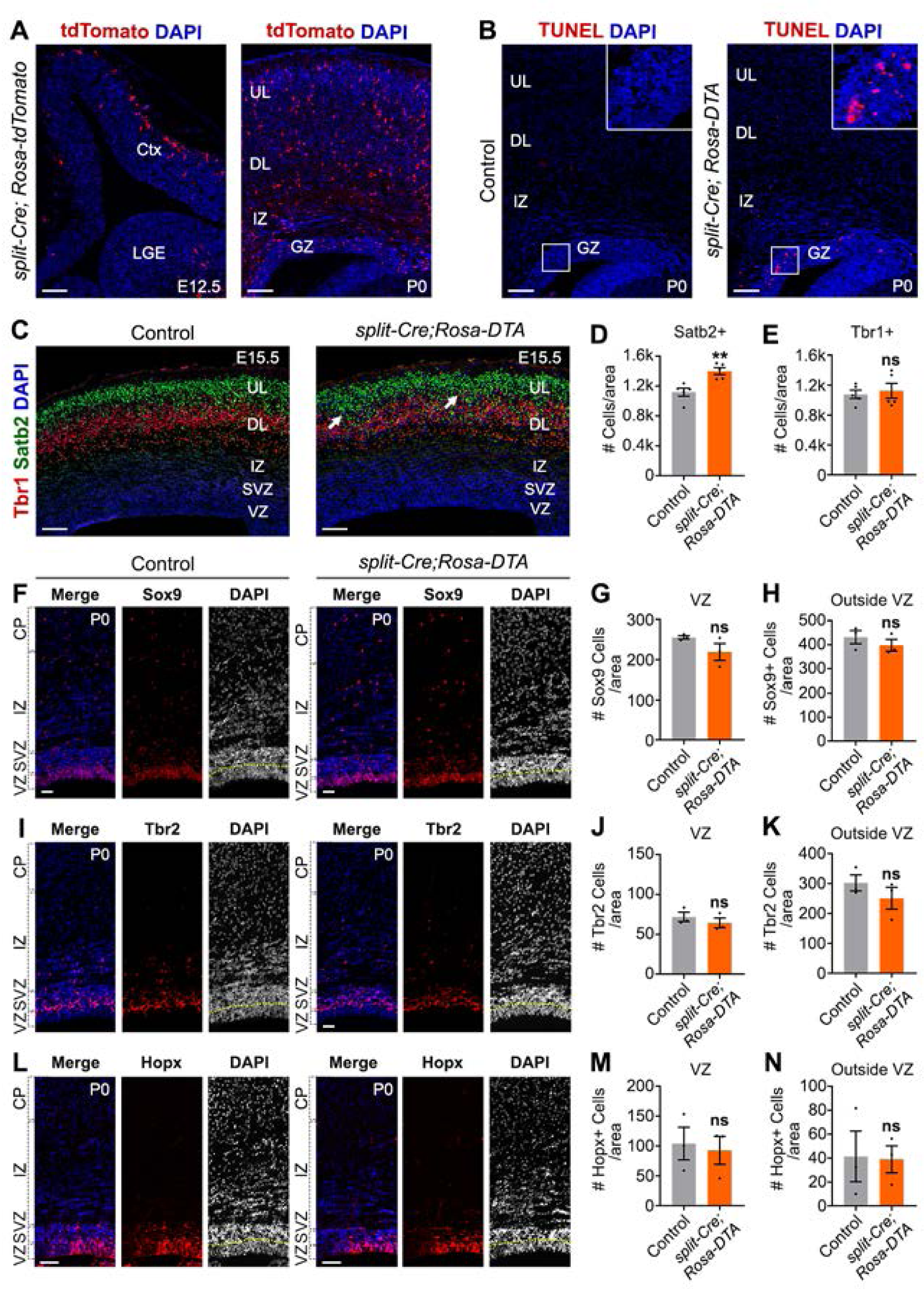

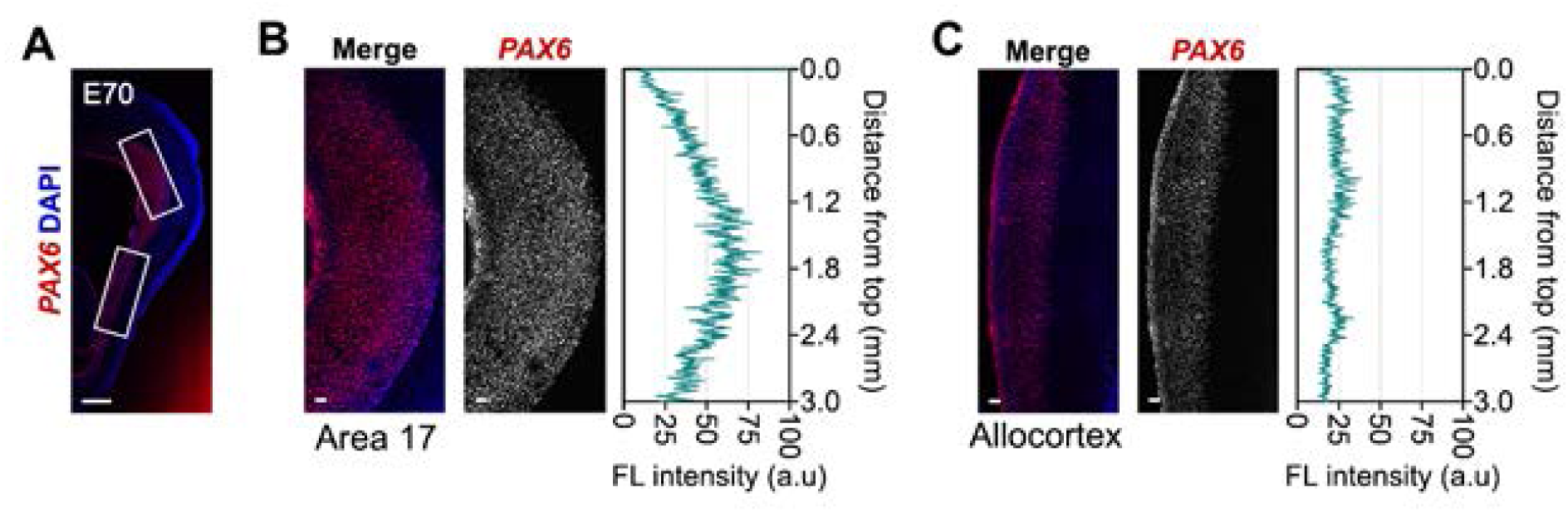

